# Bayesian Inference of Divergence Times and Feeding Evolution in Grey Mullets (Mugilidae)

**DOI:** 10.1101/019075

**Authors:** Francesco Santini, Michael R. May, Giorgio Carnevale, Brian R. Moore

**Affiliations:** Department of Evolution and Ecology, University of California, Davis, Davis, CA, U.S.A.; Dipartimento di Scienze della Terra, Universita degli Studi di Torino, Torino, Italy

## Abstract

Grey mullets (Mugilidae, Ovalentariae) are coastal fishes found in near-shore environments of tropical, subtropical, and temperate regions within marine, brackish, and freshwater habitats throughout the world. This group is noteworthy both for the highly conserved morphology of its members—which complicates species identification and delimitation—and also for the uncommon herbivorous or detritivorous diet of most mullets. In this study, we first attempt to identify the number of mullet species, and then—for the resulting species—estimate a densely sampled time-calibrated phylogeny using three mitochondrial gene regions and three fossil calibrations. Our results identify two major subgroups of mullets that diverged in the Paleocene/Early Eocene, followed by an Eocene/Oligocene radiation across both tropical and subtropical habitats. We use this phylogeny to explore the evolution of feeding preference in mullets, which indicates multiple independent origins of both herbivorous and detritivorous diets within this group. We also explore correlations between feeding preference and other variables, including body size, habitat (marine, brackish, or freshwater), and geographic distribution (tropical, subtropical, or temperate). Our analyses reveal: (1) a positive correlation between trophic index and habitat (with herbivorous and/or detritivorous species predominantly occurring in marine habitats); (2) a negative correlation between trophic index and geographic distribution (with herbivorous species occurring predominantly in subtropical and temperate regions), and; (3) a negative correlation between body size and geographic distribution (with larger species occurring predominantly in subtropical and temperate regions).

## Introduction

Grey mullets (Mugilidae, Ovalentariae) occur in coastal waters worldwide and represent an important food source in several European and Pacific countries. Mugilids are geographically widespread, with species ranging from the tropics to northern Europe, and vary greatly in body size, with species ranging from 10 − 120 cm in total length (TL). Despite this variation, the morphology of mullets is extremely conservative; all species share a torpedo-shaped body form with a similar overall appearance, which makes accurate species identification exceptionally challenging [1]. Most mullet species are euryhaline and may spend at least part of their life cycle in brackish or freshwater habitats, even though the majority of the adult life-stage and reproduction typically occur in marine habitats. However, a few species (*e.g.*, *Liza abu*, *Agonostomus monticula* and *A. catalai*) are exclusively freshwater [1–4].

The diet of grey mullets is unusual among marine fishes: most mullet species feed predominantly on food items—such as detritus and filamentous algae—with relatively low calories and/or protein per unit mass (*i.e.*, “low-quality food resources” [5]). Mullets have evolved a number of morphological adaptations associated with this diet, including a stomach with a highly muscular gizzard that serves to grind algal matter, and an extremely elongated intestine (with a variable number of pyloric caeca) that provides a greatly increased surface area to help digest and absorb algal nutrients [6]. Mugilids also possess highly modified gill rakers and a complex pharyngeal apparatus—the so-called pharyngobranchial organ [3, 7]—associated with filter feeding.

Detritivory is not uncommon in tropical freshwater habitats, having evolved independently in several distantly related freshwater lineages (*e.g.*, the characiform Prochilodontidae and Curimatidae, and several cichlid lineages [8]). By contrast, detritivory is far less common in marine groups; only ∼ 0.5% of marine fish species are predominantly herbivorous, with the vast majority of those species occurring in coral-reef habitats [9]. Recent studies suggest that the adoption of a low-quality diet may have conferred ecological opportunities that promoted rates of lineage diversification in several coral-reef groups, such as wrasses, damselfishes and surgeonfishes [5]. Surprisingly, the evolution of herbivory and detritivory in non-reef marine fishes remains largely unexplored [10].

Recent progress has greatly improved our understanding of the relationships of mugilids within acanthomorphs (ray-finned fish). Although morphological studies proposed several conflicting affinities for mugilids—including close relationships with diverse groups such as silversides and barracudas [11, 12] and sticklebacks and spiny eels [13]—a number of recent large-scale molecular studies [14–19] provide compelling evidence that mullets are members of the Ovalentaria clade [16]. Specifically, mugilids have been inferred to form a subclade with the marine surfperches (Embiotocidae) and freshwater Asiatic glassfishes (Ambassidae) [16–18].

By contrast, phylogenetic relationships *within* mugilids are far less clear; progress has been hindered by uncertainty regarding the number of species in this group, owing to the similar external morphology and widespread geographic distribution of many mullet species. Estimates for the number of mullet species peaked at 280 [2] before being reduced drastically to 75 [20]. However, recent molecular analyses suggest the possibility of many cryptic species within several widespread taxa [21]. Mullets have been the focus of intensive molecular phylogenetic analysis in the past decade [17, 21–26], which strongly contradict the traditional (morphology-based) taxonomy. Studies with the most extensive species sampling indicate that almost all non-monotypic genera are either para- or polyphyletic [21, 24]. Moreover, the number of—and relationships among—major mullet lineages remains uncertain; analyses based on multiple nuclear loci (but more limited species sampling) support three major mugilid lineages [17], whereas analyses based on three mitochondrial loci (but more intensive species sampling) support up to seven major mugilid lineages [21].

Uncertainty regarding mullet phylogeny is mirrored by uncertainty regarding a time scale for their diversification. Previous estimates of divergence times in this group have included a limited number of extant mullet species and/or fossil calibrations. For example, a large-scale study of vertebrate divergence times included 24 mullet species and no mullet fossils [19], a large-scale study of teleost divergence times included 10 mullet species and no mullet fossils [17], a large-scale study of acanthomorph divergence times included 2 mullet species and no mullet fossil calibrations [18], and a smaller-scale study included seven extant mullet species and a single mullet fossil calibration [25].

Here, our main objectives are to provide a comprehensive estimate of the phylogenetic relationships and divergence times within mullets, and to use the resulting phylogeny as a framework to explore the evolution of feeding preference in this group. To this end, we perform a series of statistical analyses to: (1) estimate the number of distinct mullet species; (2) estimate the phylogeny and divergence times for the identified species; (3) infer the evolutionary history of feeding preference using the resulting dated phylogeny, and; (4) explore correlations between feeding preference and several other variables, including body size, habitat (marine, brackish or freshwater), and geographic distribution (tropical, sub-tropical, or temperate).

## Materials and Methods

### Sequence data

We obtained 282 mullet sequences from GenBank for three mitochondrial genes: 16S, COI, and cyt*b*. We first excluded 19 sequences that were identified only to the generic level: *Chelon* sp. (1 sequence), *Liza* sp. (9 sequences), *Moolgarda* sp. (4 sequences), and *Valamugil* sp. (5 sequences). We excluded 32 additional sequences with 100 percent sequence identity (phylogenetic models assume a binary tree topology, which is violated by datasets that include multiple identical sequences). Finally, we included two embiotocid (surfperch) species as outgroups: *Cymatogaster aggregata* and *Ditrema temninckii*.

We aligned sequences for each gene using `MUSCLE` v.3.8.31 [27], confirmed the reading frame by examining the amino-acid translation in `AliView` v.1.18 [28], and then trimmed the ragged 3*′* and 5*′* ends of each aligned gene. The concatenated alignment comprised a total of 1986 sites—including 604 bp of 16S, 598 bp of COI, and 784 bp of cyt*b*—for a total of 233 sequences, with 5.4% missing data (Table S9).

### Comparative data

For every species in our study, we scored several discrete and continuous variables (Table S10), including: (1) feeding preference (FP), expressed as a discrete variable with three states (algae, detritus, or invertebrates); (2) total body length (TL), expressed as a continuous variable in centimeters; (3) trophic index (TI), expressed as a continuous variable based on stomach contents; (4) habitat type (Hab.), expressed as a discrete variable with three states—marine or non-marine (brackish or freshwater) — reflecting the environment in which each species spends most of its life cycle; (5) geographic distribution (Dist.), expressed as a discrete variable with three states—tropical or non-tropical (subtropical or temperate). We gathered these data from various sources, including FishBase [20], the FAO fish identification guides [3], and the survey of geographic distribution by Briggs and Bowen [29].

#### Data Availability Statement

The authors confirm that all data supporting the results of this study are fully available without restriction. These data are available as an archive—including all molecular and comparative data (and the corresponding input files with full model specification in NEXUS and XML formats)—deposited in the Dryad database. The Dryad data identifier is: doi:10.5061/dryad.h26v3 (viewable at http://datadryad.org/review?doi=doi:10.5061/dryad.h26v3).

### Species delimitation

We first sought to estimate the number of distinct mullet species within the 233 sequence dataset using the Poisson tree process (PTP) model [30]. This approach requires a single, rooted phylogram as input (*i.e.*, with branch lengths rendered as the expected number of substitutions per site). To this end, we estimated the phylogeny for the 233 sequence dataset. We selected a mixed substitution model (partition scheme) for this dataset using `PartitionFinder` v.1.1.1 [31]. We defined 8 data subsets—one for each codon position of the two protein-coding genes, and one each for the stem and loop regions of the 16S ribosomal gene—and explored the space of partition schemes using the heuristic (‘greedy’) algorithm to search among the set of substitution models implemented in `MrBayes` v. 3.2.4 [32], and used the Bayesian Information Criterion (BIC) [33] to select among the candidate partition schemes (Table S1).

We then estimated the posterior probability distribution of trees (and other model parameters) under the selected mixed substitution model. Specifically, we approximated the joint posterior probability distribution using the Markov chain Monte Carlo (MCMC) algorithms implemented in `MrBayes` v.3.2.4 [32], running six independent, replicate simulations for 10^8^ cycles, and thinned each chain by sampling every 10,000^th^ state. To assess the reliability of the MCMC simulations, we used the `Tracer` [34] and `coda` [35] packages. Namely, we assessed convergence of each MCMC simulation to the stationary (joint posterior) distribution by plotting the time series for every parameter, and calculated both the effective sample size (ESS) [36] and Geweke (GD) [37] diagnostics for every parameter. We assessed mixing of each chain over the stationary distribution by calculating both the potential scale reduction factor (PSRF) [38] diagnostic and monitoring the acceptance rates for all parameters. Additionally, we assessed convergence of the MCMC simulations by comparing the six independent estimates of the marginal posterior probability density for each parameter, ensuring that all parameter estimates were effectively identical and SAE compliant [36]. Based on these diagnostic analyses, we discarded the first 50% of samples from each chain as burn-in, and based parameter estimates on the combined stationary samples from each of the six independent chains (*N* = 30, 000). We summarized the resulting composite marginal posterior distribution of phylogenies as an all-compatible majority-rule consensus tree, and rooted the consensus tree using the two outgroup species (Figures S1–S2).

The resulting rooted phylogram served as the (pseudo)data for delimiting mullet species. We performed Bayesian inference under the PTP model using the stand-alone implementation of `bPTP` [30], running four replicate MCMC simulations for 20 million cycles, sampling every 2,000^th^ state, and assessed the reliability of the simulations as described above. The resulting set of 100 distinct species (comprising 98 mullet and two surfperch species; Figures S2–S3) were used for all subsequent statistical analyses (to infer the phylogeny, divergence times, and evolution of feeding preference in mullets).

### Phylogeny and divergence-time estimation

We inferred divergence times within a Bayesian statistical framework using relaxed-clock models. These models comprise three main components [39]: (1) a *site* model describes how the nucleotide sequences evolved over the tree with branch lengths, while accommodating variation both in the *rate* of substitution across sites and the *nature* of the substitution process across sites (*i.e.*, by means of ‘partition schemes’); (2) a *branch-rate* prior model specifies how substitution rates are distributed among branches of the phylogeny, and; (3) a *node-age* prior model specifies the distribution of speciation times in the phylogeny. Additionally, estimating *absolute* divergence times requires the inclusion of one or more *calibrations* to scale relative ages to absolute, geological time. To estimate divergence times in mullets, we evaluated the fit of our sequence data to six candidate relaxed-clock models—comprising all combinations of two site models, three branch-rate models, and one node-age model—and used three fossil calibrations.

#### Relaxed-clock models

We selected site models for the 100-species dataset using `PartitionFinder` v.1.1.1 [31]. Specifically, we used the heuristic (‘greedy’) algorithm to explore the space of partition schemes for the set of substitution models implemented in `BEAST` v. 1.8.2 [34], and selected among the candidate partition schemes using both the Bayesian Information Criterion (BIC) [33] and the Akaike Information Criterion (AIC) [40]. The two resulting partition schemes—PS1 selected using the BIC, and PS2 selected using the AIC—are summarized in Table 1.

**Table 1.** Mixed-model selection. We selected among the space of partition schemes that variously assign substitution models to 8 data subsets using both the BIC and AIC model-selection methods implemented in `PartitionFinder`. The number of free substitution-model parameters (excluding branch lengths) for each of the partition schemes is indicated in the rightmost column, NP.

| Partition | Data Subset |  |  |  |  |  |  |  |  |
| --- | --- | --- | --- | --- | --- | --- | --- | --- | --- |
|  | *cox1* |  |  | *cytb* |  |  | 16S |  |  |
|  | 1<sup>st</sup> pos. | 2<sup>nd</sup> pos. | 3<sup>rd</sup> pos. | 1<sup>st</sup> pos. | 2<sup>nd</sup> pos. | 3<sup>rd</sup> pos. | stem | loop | NP |
| PS1(BIC) | K80+$\Gamma$ | HKY+I | TrN+$\Gamma$ | SYM+$\Gamma$ | TrN+$\Gamma$ | GTR+$\Gamma$ | SYM+$\Gamma$ | GTR+$\Gamma$ | 62 |
| PS2(AIC) | TrN+$\Gamma$ | TrN+$\Gamma$ | GTR+$\Gamma$ | SYM+$\Gamma$ | TrN+$\Gamma$ | GTR+$\Gamma$ | SYM+$\Gamma$ | GTR+$\Gamma$ | 72 |

For both of the selected partition schemes (PS1 and PS2), we evaluated three branch-rate models to describe how substitution rates vary across branches of the tree. Specifically, we evaluated: (1) the uncorrelated lognormal (UCLN) [41] model, which assumes that substitution rates on adjacent branches are drawn from a shared lognormal distribution; (2) the uncorrelated exponential (UCEX) [41] model, which assumes that substitution rates on adjacent branches are sampled from a shared exponential distribution, and; (3) the random-local clock (RLC) [42] model, which assumes that substitution rates are locally constant within sections of the tree, where the number and distribution of constant-rate sections are modeled as a truncated Poisson process.

To complete the relaxed-clock model specification, we chose the sampled birth-death (SBD) branching process model [43] to describe the prior distribution of branching times in the tree, as it accommodates both extinction and incomplete species sampling. Other potential node-age prior models were discounted *a priori* on biological grounds. For example, the pure-birth (Yule) [44] branching process model—which assumes a zero extinction rate—is inappropriate in light of fossil evidence documenting extinction in mullets. Similarly, the birth-death (BD) [45] branching process model—which assumes complete species sampling—is violated by the incomplete (albeit comprehensive) species sampling used here.

#### Fossil calibrations

In order to estimate *absolute* divergence times, we applied fossil calibrations as prior probability densities to three internal nodes of the mullet phylogeny. Because fossil calibrations are typically applied by constraining the monophyly of the corresponding internal nodes, we first performed a series of preliminary analyses (under each of the candidate relaxed-clock models) to estimate the posterior probability for each internal node that represented a prospective calibration point. These analyses inferred strong support for the three prospective calibration points (Table 2); accordingly, we constrained each of the calibrated nodes to be monophyletic when estimating divergence times. We assigned calibrations to internal nodes of the phylogeny—and specified the form (hyperpriors) of the corresponding prior probability densities—based on the morphological features of the fossils, and on the stratigraphy of the horizons from which the fossils are known, respectively. We discuss these considerations for each of the fossil calibrations below.

**Table 2.**
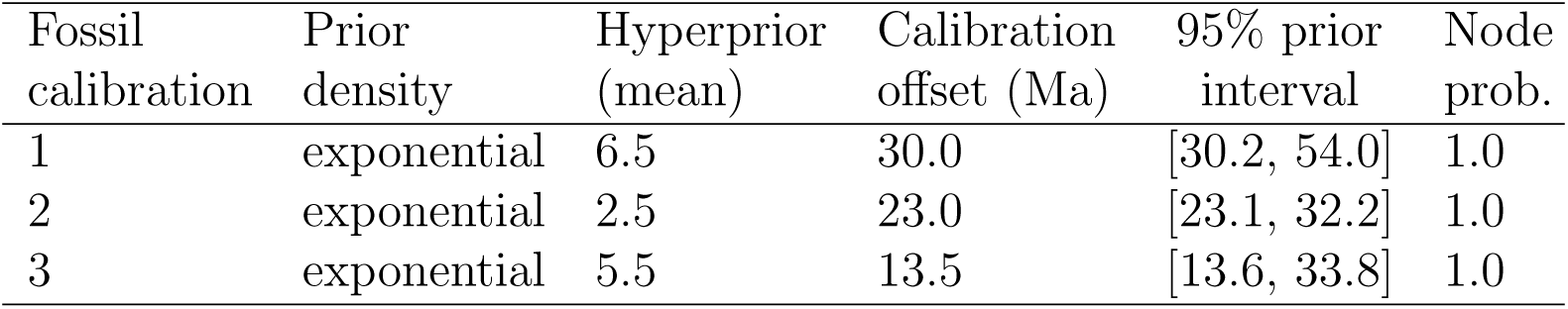
Prior probability densities for fossil calibrations. Numbers for the three fossil calibrations correspond both to those used in the text, and to the indices of the internal nodes on the trees in Figures 1 and S7. The posterior probability for each calibrated node—estimated without topological constraints imposed—is indicated in the rightmost column.

| Fossil calibration | Prior density | Hyperprior (mean) | Calibration offset (Ma) | 95% prior interval | Node prob. |
| --- | --- | --- | --- | --- | --- |
| 1 | exponential | 6.5 | 30.0 | [30.2, 54.0] | 1.0 |
| 2 | exponential | 2.5 | 23.0 | [23.1, 32.2] | 1.0 |
| 3 | exponential | 5.5 | 13.5 | [13.6, 33.8] | 1.0 |

##### Fossil calibration 1

Articulated skeletal remains of *Mugil princeps* from the Menilite Shales of Ukraine [46] document the presence of the genus *Mugil* in the Rupelian, 34 − 28 million years ago (Ma). Our inclusion of this fossil within the crown *Mugil* clade is supported by its possession of a maxilla with a straight posterior end, absence of an opercular spine, and arrangement and meristics of median fins [2, 47, 48]. The age of the Menilite-type shales of the Outer Carpathians has been studied extensively [49], which refer these fossiliferous deposits to the NP23 nanoplankton zone that is estimated to have a minimum age of ∼ 30 Ma. *Mugil princeps* has also been reported from the slightly younger Rupelian Menilites Shales of Poland and the Chattian brackish deposits of Aix-en-Provence in southern France [46, 50]. A single otolith belonging to an unspecified mullet (Mugilidae *indet.*) from the Santonian “Arcillas et Margas de la Font de las Bagasses”, in Catalonia, Spain [51] constitutes the earliest mullet fossil remains. The “Arcillas et Margas de la Font de las Bagasses” belongs to the *Dicarinella asymetrica* planktonic foraminifera zone; the Late Santonian age (84.5 − 83.5 Ma) of these deposits is also supported by the presence of the ammonite *Placenticeras syrtale* [52, 53]. This Santonian otolith reflects a probable upper bound on the age this node, which has a minimum age of 30 Ma (Table 2).

##### Fossil calibration 2

The otolith-based species *Chelon gibbosus* from the Chattian brackish deposits of the Gr′es et Marnes gris à gypse Formation (=Untere Suüsswassermolasse), in the western part of Switzerland, provides a minimum age for the clade comprising *Chelon labrosus*, *Liza aurata*, *Liza dumerili*, *Liza saliens*, *Liza richardsoni*, *Liza bandaliensis* and *Liza tricuspidens*. Reichenbacher and Weidman [54] demonstrated remarkable similarities between this Oligo-Miocene taxon and the extant *Chelon labrosus*. The fossiliferous layers of the Gr′es et Marnes gris à gypse Formation are stratigraphically referred to the MP30 mammal zone [54], with a minimum age of ∼ 23 Ma [55]. Accordingly, the corresponding calibration is specified with a minimum age of 23 Ma (Table 2).

##### Fossil calibration 3

Otoliths referred to *Mugil* aff. *cephalus* from the Miocene Cantaure Formation, Paraguana Peninsula, of Venezuela [56], provide a minimum age for the extant species *Mugil cephalus*. The Miocene otoliths from the Cantaure Formation are identical to those of Recent individuals of the flathead grey mullet *Mugil cephalus* [56]; however, the juvenile nature of these Miocene otoliths renders their identification uncertain. The age of the Cantaure Formation has been carefully studied [57]; these deposits have been assigned to the Burdigalian-Langhian NN4 and NN5 nanoplankton zones, which have a minimum age of ∼ 13.65 Ma [58]. Accordingly, we specified the corresponding calibration with a minimum age of 13.65 Ma, and a soft upper bound of 30 Ma (Table 2).

#### Relaxed-clock model selection

We evaluated the fit of the sequence data to the six candidate relaxed-clock models—including all combinations of the two partition schemes (PS1, PS2), three branch-rate models (UCLN, UCED, RLC), and single node-age model (SBD)—using robust Bayesian model-selection methods. This *Bayes factor* approach involves first estimating the *average* fit of the data to each candidate model—where the likelihood of the data is averaged over the joint prior probability density of the model parameters (the *marginal likelihood*)—and then assessing the relative fit of the competing models by comparing their marginal-likelihood values [59].

We estimated the marginal likelihood of each candidate model using robust (albeit computationally intensive) ‘stepping-stone’ [60, 61] and ‘path-sampling’ estimators [62, 63]. These algorithms are similar to the familiar MCMC algorithms, which are intended to sample from (and estimate) the joint posterior probability of the model parameters. Stepping-stone algorithms are like a series of MCMC simulations that iteratively sample from a specified number of discrete steps between the posterior and the prior probability distributions. The basic idea is to estimate the probability of the data for all points between the posterior and the prior—effectively summing the probability of the data over the prior probability of the parameters to estimate the marginal likelihood.

We estimated the marginal likelihood for each of the candidate relaxed-clock models—using proper priors for all parameters—by simulating from the posterior to the prior across 100 stones. We ran each simulation for a total of 1 billion cycles, visiting each stone for 10^7^ cycles, thinning the chain by sampling every 1000^th^ state, and discarding the first 10% of samples from each stone. We distributed the stones between the posterior and prior as evenly spaced quantiles of a beta distribution, with the shape parameters specified to concentrate stones near the prior, Beta(0.3, 1.0). To assess the stability of the marginal likelihood estimates, we performed three replicate stepping-stone simulations for each of the six candidate relaxed-clock models. Finally, we used the resulting marginal likelihood estimates (Table 3) to select among the corresponding relaxed-clock models using Bayes factors (Table 4).

**Table 3.**
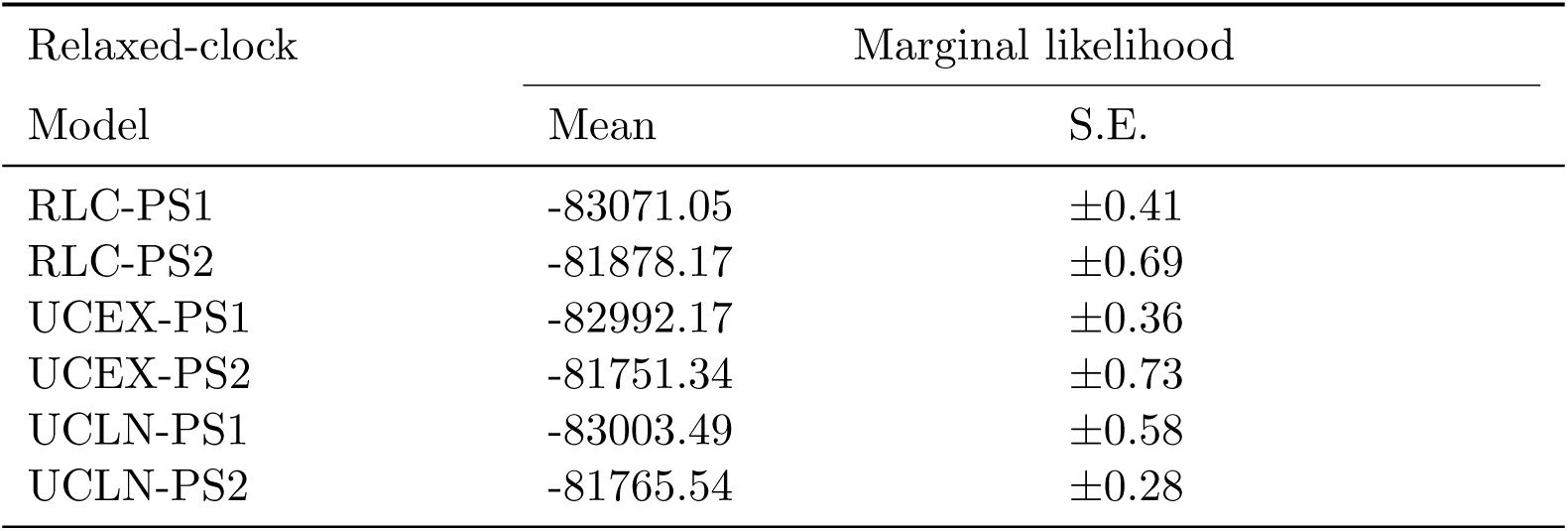
Marginal-likelihood estimates for relaxed-clock models. Model comparisons are based on analyses of the 100-species dataset. Marginal likelihoods for each of the candidate relaxed-clock models are based on the stepping-stone estimator [63, 64]. Estimates of the standard error (S.E.) are based on 1000 bootstrap replicates performed in `Tracer` v.1.6.

**Table 4.** Bayes factor comparisons for relaxed-clock models. Model comparisons are based on analyses of the 100-species dataset. For each model comparison, *M*_0_ : *M*_1_, we calculated the Bayes factor as 2*ln*(*M*_0_ − *M*_1_). The table compares marginal likelihoods for the pair of models in row *i* and column *j*: positive values indicate support for the model in row *i*. The UCEX-PS2-SBD relaxed-clock model is decisively preferred over rival models (*ln*BF > 4.6) [65].

| Relaxed-clock<br>Model | Bayes Factor |  |  |  |  |  |
| --- | --- | --- | --- | --- | --- | --- |
|  | RLC-PS1 | RLC-PS2 | UCEX-PS1 | UCEX-PS2 | UCLN-PS1 | UCLN-PS2 |
| RLC-PS1 | — | -1192.88 | -78.87 | -1319.71 | -67.56 | -1305.51 |
| RLC-PS2 | 1192.88 | — | 1114.01 | -126.82 | 1125.38 | -112.62 |
| UCEX-PS1 | 78.87 | -1114.01 | — | -1240.83 | 11.32 | -1226.63 |
| UCEX-PS2 | 1319.71 | 126.82 | 1240.83 | — | 1252.15 | 14.20 |
| UCLN-PS1 | 67.56 | -1125.33 | -11.32 | -1252.15 | — | -1237.95 |
| UCLN-PS2 | 1305.51 | 112.62 | 1226.63 | -14.20 | 1237.95 | — |

#### Parameter estimation

We estimated the joint posterior probability distribution of the phylogeny, divergence times and other parameters under the selected relaxed-clock model—the UCEX-PS2-SBD model—and three fossil calibrations using the MCMC algorithms implemented in `BEAST` v.1.8.2 [34]. Specifically, we ran four replicate MCMC simulations for 10^8^ cycles, thinned chains by sampling every 10, 000^th^ state, and assessed the reliability of the approximations as described previously. We then combined the stationary samples from the four independent simulations, and summarized the resulting composite marginal posterior probability density as a maximum clade credible (MCC) consensus tree with median node ages (Figure 1).

**Figure 1.**
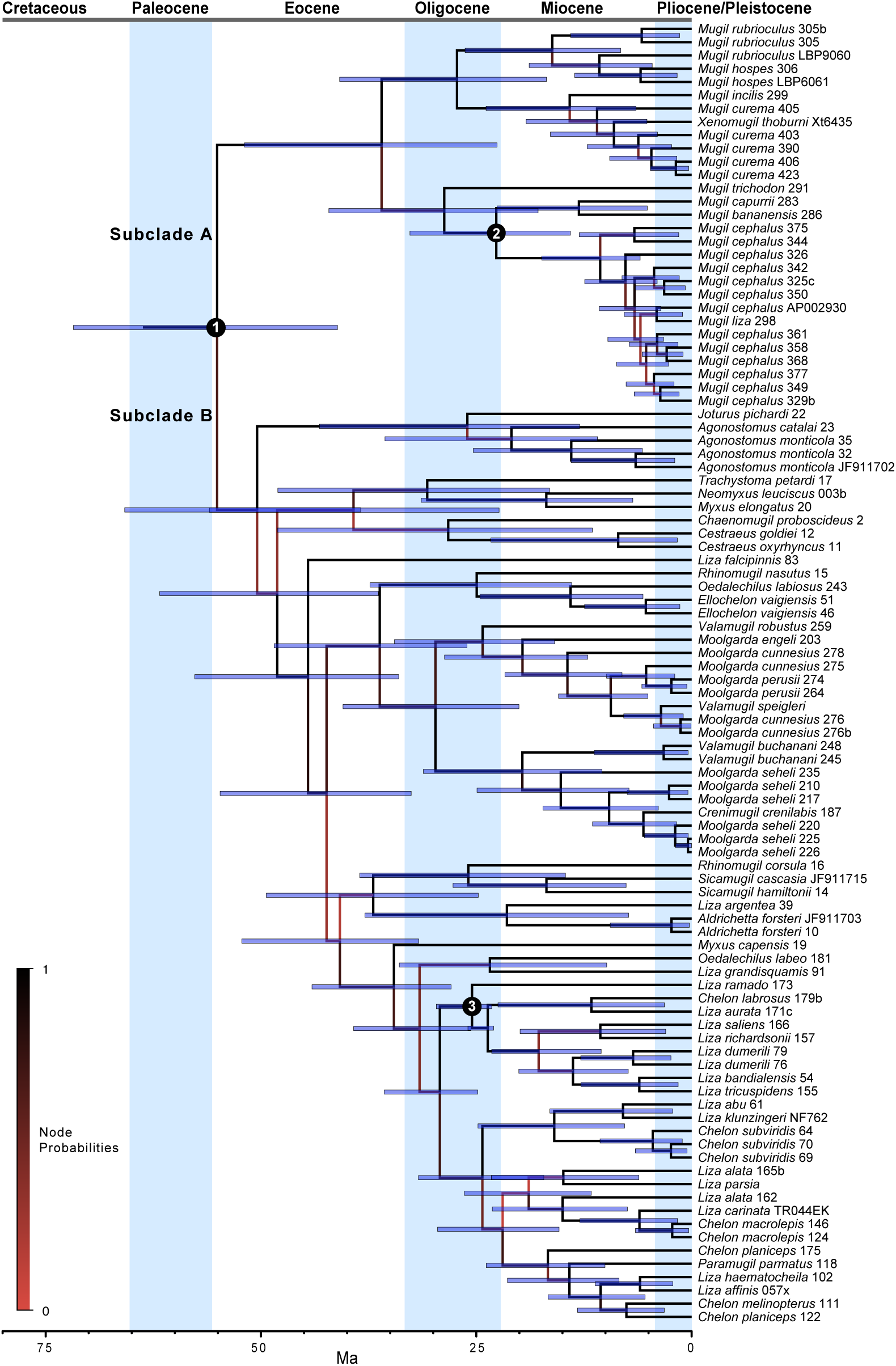
Bayesian estimate of mullet phylogeny and divergence times. The shading of internal branches indicates the corresponding node probabilities (see inset legend), the numbered internal nodes indicate the location of the corresponding fossil calibrations (see Table 2), and the bar plots on nodes indicate the corresponding 95% HPD interval of divergence times.

### Ancestral-state estimation

We used the inferred phylogeny as a framework for exploring the evolution of feeding preference in mullets. We scored feeding preference as a discrete variable with three states (algae, detritus, or invertebrates), reflecting the main food item of each mullet species. We assessed the fit of these discrete traits to six candidate models, comprising all possible combinations of two continuous-time Markov (CTM) models (that describe the instantaneous rates of change between the discrete states) and three branch-rate models (that describe how rates of diet evolution vary across branches of the tree). Specifically, we evaluated one trait models that assumes a single, symmetric instantaneous rate of change between each pair of states (CTM-3), and a second model that assumes independent, asymmetric rates of change between all states (CTM-6). The three branch-rate models include the constant-rate morphological clock model (CRMC), where the rate of trait evolution is assumed to be constant across branches, and the uncorrelated exponential model (UCEX), where rates of trait evolution and substitution vary across branches under a *shared* branch-rate model (UCEX_*s*_), or where rates of trait evolution and substitution vary across branches under *independent* branch-rate models (UCEX_*i*_).

We conditioned inferences of diet evolution on the previously inferred MCC topology (Figure 1), but integrated out uncertainty in divergence times under the preferred relaxed-clock model (UCEX-PS2-SBD; Table 4) and the three fossil calibration densities (Table 2). We assumed uniform priors, Uniform(0, 1), for both the stationary and root frequencies of the three discrete states, and a mean-one gamma prior, Gamma(1, 1), on the instantaneous-rate parameters. We simultaneously estimated the number of changes in feeding preference in mullets—between diets of algae, detritus, or invertebrates—using the robust Markov-jump approach [66, 67] implemented in `BEAST` v.1.8.2 [34].

For each candidate discrete-trait model, we inferred the joint posterior probability by performing four replicate MCMC simulations of 400 million cycles in `BEAST` v.1.8.2 [34], thinning the chain by sampling every 4000^th^, and assessed the reliability of the approximations as previously. We combined the stationary samples from the four replicate simulations under each model, and used these composite posterior samples to assess the fit of the discrete-trait data to the the four candidate models. Specifically, we estimated the marginal likelihood for each discrete-trait model using the AICm method-of-moments estimator [63, 64, 68] implemented in `Tracer` v.1.6 [34]. We then used the resulting marginal likelihood estimates (Table 5) to select among the corresponding discrete-trait models using Bayes factors (Table 6). Finally, we plotted the marginal probabilities for diet on the internal nodes of the MCC consensus tree using `FigTree` v.1.4.2 (Figure 2), and summarized the instantaneous rates and number of changes between states (Table 9).

**Figure 2.**
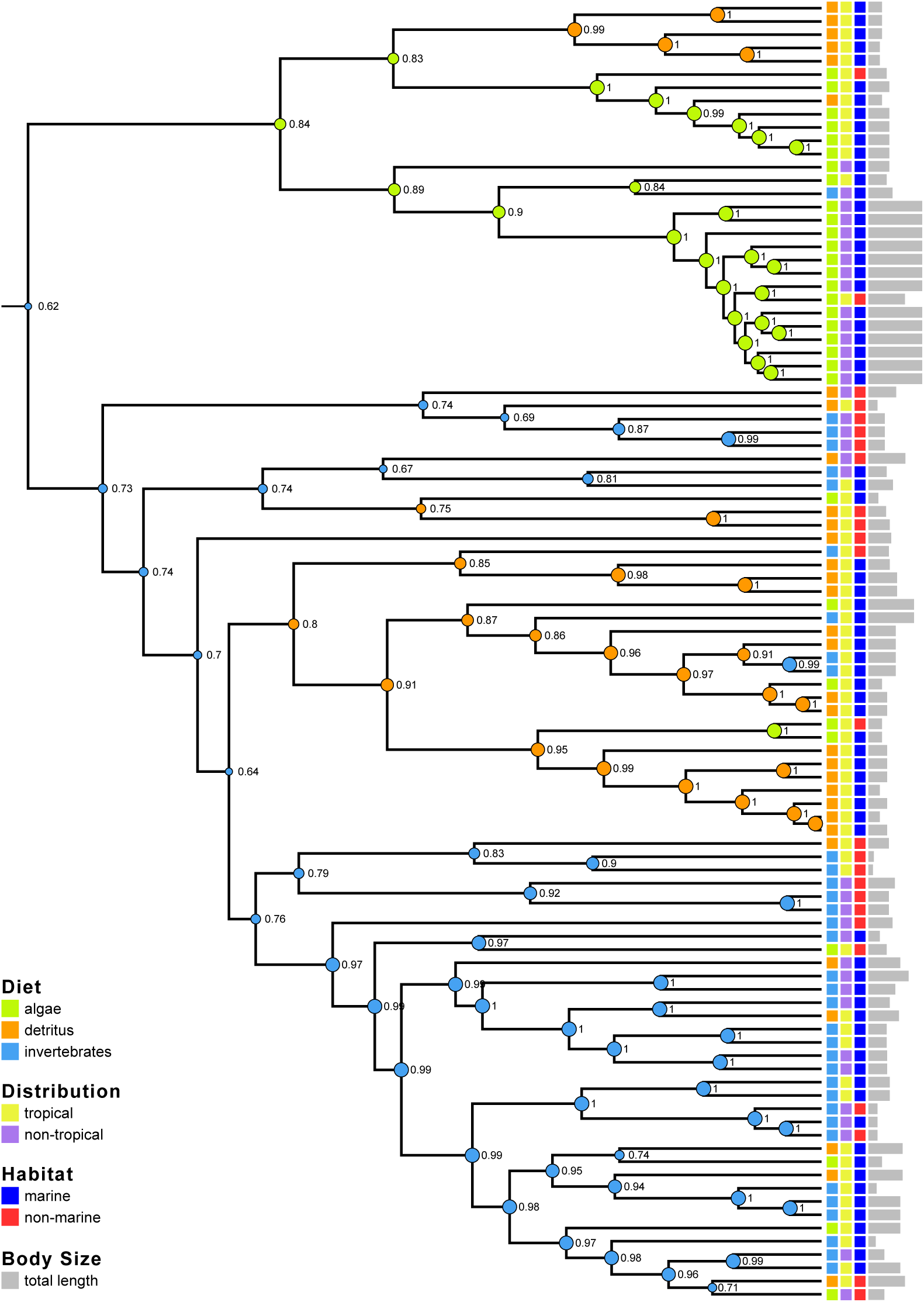
Bayesian inference of diet evolution in mullets. Circles at interior nodes are colored according to the MAP estimate of the ancestral diet—algae, detritus, or invertegrates—here the diameter and adjacent numbers indicate the marginal posterior probability of the MAP state. Other variables—biogeographic distribution, habitat, and body size—are indicated at the tips of the tree for each species (see inset legend).

**Table 5.** Marginal-likelihood estimates for discrete-trait models. Model comparisons are based on analyses of the 100-species dataset. Candidate discrete-trait models comprise all combinations of branch-rate models—the constant-rate morphological clock (CRMC) model and the uncorrelated-exponential relaxed-clock (UCEX) models, where rates of substitution and diet evolution are either shared (*s*) or independent (*i*)—and site models—where rates of change between the three discrete-traits are assumed to be symmetric (CTM3) or are allowed to be asymmetric (CTM6). Marginal likelihoods for each of the candidate discrete-trait models are based on the AICm method-of-moments estimator [63, 64]. Estimates of the standard error (S.E.) are based on 1000 bootstrap replicates performed in `Tracer` v.1.6.

| Discrete-trait<br>Model | Marginal likelihood |  |
| --- | --- | --- |
|  | Mean | S.E. |
| CRMC-CTM3 | -81841.34 | ±0.63 |
| CRMC-CTM6 | -81845.93 | ±0.59 |
| UCEX<sub>s</sub>-CTM3 | -81818.36 | ±0.64 |
| UCEX<sub>s</sub>-CTM6 | -81786.74 | ±0.60 |
| UCEX<sub>i</sub>-CTM3 | -81844.71 | ±0.53 |
| UCEX<sub>i</sub>-CTM6 | -81860.50 | ±0.70 |

**Table 6.** Bayes factor comparisons for discrete-trait models. Model comparisons are based on analyses of the 100-species dataset. For each model comparison, *M*_0_ : *M*_1_, we calculated the Bayes factor as 2*ln*(*M*_0_ *− M*_1_). The table compares marginal likelihoods for the pair of models in row *i* and column *j*: positive values indicate support for the corresponding model in row *i*. The shared UCEX-CTM 6-rate discrete-trait model is decisively preferred over rival models (*ln*BF > 4.6) [65].

| Discrete-trait<br>Model | Bayes Factor |  |  |  |  |  |
| --- | --- | --- | --- | --- | --- | --- |
|  | CRMC-3 | CRMC-6 | UCEX<sub>s</sub>-3 | UCEX<sub>s</sub>-6 | UCEX<sub>i</sub>-3 | UCEX<sub>i</sub>-6 |
| CRMC-CTM3 | — | 4.59 | -22.98 | -54.60 | 3.37 | 19.16 |
| CRMC-CTM6 | -4.59 | — | -27.57 | -59.19 | -1.22 | 14.58 |
| UCEX<sub>s</sub>-CTM3 | 22.98 | 27.57 | — | -31.62 | 26.35 | 42.15 |
| UCEX<sub>s</sub>-CTM6 | 54.60 | 59.19 | 31.62 | — | 57.97 | 73.77 |
| UCEX<sub>i</sub>-CTM3 | -3.37 | 1.22 | -26.35 | -57.97 | — | 15.80 |
| UCEX<sub>i</sub>-CTM6 | -19.16 | -14.58 | -42.15 | -73.77 | -15.80 | — |

### Correlated-trait evolution

We explored correlations among traits using the recently developed multivariate phylogenetic latent-liability model [69]. Briefly, this method estimates pairwise correlation coefficients among a set of discrete and continuous traits by treating the discrete trait values for each species as “latent” (unobserved) continuous traits. The combined continuous and latent traits are assumed to evolve under a correlated Brownian motion model with variance-covariance matrix, **∑**, which is a square matrix with a number of rows and columns equal to the number of traits being studied.

The elements of **∑** contain the parameters of interest: the diagonal elements, **∑**_*ii*_, represent the evolutionary rate of trait *i*, while the off-diagonal elements, **∑**_*ij*_, represent the covariance between traits *i* and *j*. These parameters are estimated in a Bayesian statistical framework; it is therefore necessary to specify prior values for the precision matrix, **∑**^−1^, and hyperparameters for the rate matrix, **R**, and degrees of freedom, ***d***.

We assessed correlations among four continuous and discrete traits: trophic index (TI, continuous), total length (TL, continuous), habitat (Hab, discrete), and distribution (Dist, discrete). The latent-liability model assumes continuous traits can realize any positive or negative value. Accordingly, it was necessary to transform our continuous traits to satisfy this assumption. Trophic index (which ranges from 2.0 to 3.4) was normalized (to range from 0 and 1), and subsequently logit-transformed. The logit-transformed trophic index values range between −∞ to ∞. Total length (which ranges from 0 to *∞*) was ln-transformed, resulting in values between −∞ and ∞. We treated both discrete traits as binary; habitat was scored as marine and non-marine, while distribution was scored as tropical and non-tropical.

We explored correlated-trait evolution in mullets on the MCC tree (Figure 1) using the latent-liability model implemented in `BEAST` v. 1.8.3. To assess the sensitivity of inferred trait correlations to the choice of priors, we explored three different values for the rate-matrix prior, **R** (low, medium, and high), and three different values for the precision-matrix prior, **∑**^−1^ (low, medium, and high). We chose to use a fixed value of ***d*** = 6 for all analyses (*i.e.*, the number of traits plus two). For each combination of prior settings (9 in total), we ran four independent MCMC simulations for 200 million cycles, thinning each by sampling every 20,000^th^ state, providing 10,000 samples per simulation. We assessed performance of the MCMC simulations in the usual manner. We then combined the stationary samples from the four independent simulations for each of the 9 prior combinations; the resulting composite marginal posterior probability densities were used to estimate the marginal posterior densities of covariances among traits.

Finally, we transformed the marginal densities of evolutionary covariances into marginal densities of correlation coefficients, which range from −1 to 1 to provide a more natural interpretation of correlations that can be compared among traits, regardless of the overall rate of evolution. For each marginal density, we identified the correlation coefficient as significantly different from zero if a correlation coefficient of zero (*i.e.*, no correlation) was as or more extreme than 95% of the marginal density.

### Sensitivity analyses

The analyses described above—to estimate the phylogeny and divergence times of mullets, and to explore the evolution of their feeding preference using this dated tree—are based on the set of 98 delimited mullet species. Given the historical difficulties in defining the number of species within this morphologically conservative group, and potential limitations of our attempt to objectively delimit species from all available mitochondrial sequence data, we sought to assess the sensitivity of our findings to uncertainty in the delimitation of mullet species.

To this end, we defined a dataset comprising the conventionally recognized mullet species by randomly selecting a single sequence for each of the 62 nominal mullet species represented in the 233-sequence dataset. That is, for every species with *N >* 1 sequences, we randomly selected a single sequence (where each sequence was selected with a probability of 1*/N*) without reference to the phylogenetic position of the sequences or the values of other variables (diet, body length, habitat, or geographic distribution).

We then repeated the entire series of analyses described above for the 98 mullet species and two outgroup species (*i.e.*, the ‘100-species dataset’) for the 62 mullet and two outgroup species (*i.e.*, the ‘64-species dataset’). These analyses and results are described in the Supporting Information (Tables S2–S8; Figures S6–S9). Overall, our study entailed approximately 500 analyses that consumed ∼ 66, 000 hours (∼ 7.6 years) of compute time. All of the analyses for this study we performed on the CIPRES Science Gateway v.3.3 [70].

## Results and Discussion

### Species delimitation in mullets

Our species-delimitation analyses of the 233-sequence dataset identified 98 distinct mullet species (adding 36 novel species to the 62 recognized mullet species represented in the 233-sequence dataset; Figure S3). However, we emphasize that we do not view these results as definitive. First, our analyses are based exclusively on mitochondrial gene regions, which raises concerns about possible confounding effects of introgression. Second, the scale of the mullet dataset required use of relatively efficient (but approximate) species-delimitation methods (based on the Poisson tree process model [30]), which provide an approximation of more theoretically sound methods (based on multi-species coalescence models [71–74]). Unfortunately, these more rigorous species-delimitation approaches were not computationally viable for the mullet dataset. Finally, our results are (necessarily) based on a finite sample of individuals and gene regions. Our inferences regarding the number of distinct mullet species would likely change if we were to: (1) increase the sample of individuals for the same mitochondrial genes; (2) increase the geographic scope of sampled individuals for the same mitochondrial genes, and/or; (3) increase the scope of gene/omic regions for the same individuals.

In light of the historical difficulties in delimiting species within this morphologically conservative group, we fully anticipate that the number of recognized mullet species will change as the geographic and genomic sampling of this group continues to improve. Nevertheless, our estimates are presented as an attempt to objectively quantify the number of distinct mullet species based on the most comprehensive sample of molecular sequence data currently available. Moreover, our findings regarding the newly delimited mullet species at least seem biologically plausible in light of other lines of independent evidence. Specifically, most (33 of 36) of the newly delimited species correspond to geographically isolated species clusters identified by Durand and colleagues [21, 24]. For example, we identified two distinct species from geographically isolated clusters of *Moolgardia perusi*, three distinct species from geographically isolated clusters of *Agonostomus monticola*, four distinct species from isolated clusters of *Moolgardia cunnesius*, five distinct species from isolated clusters of *Mugil curema*, six distinct species from isolated clusters of *Moolgardia seheli*, and 13 distinct species within the circumglobal *Mugil cephalus* species complex (Table 7).

**Table 7.**
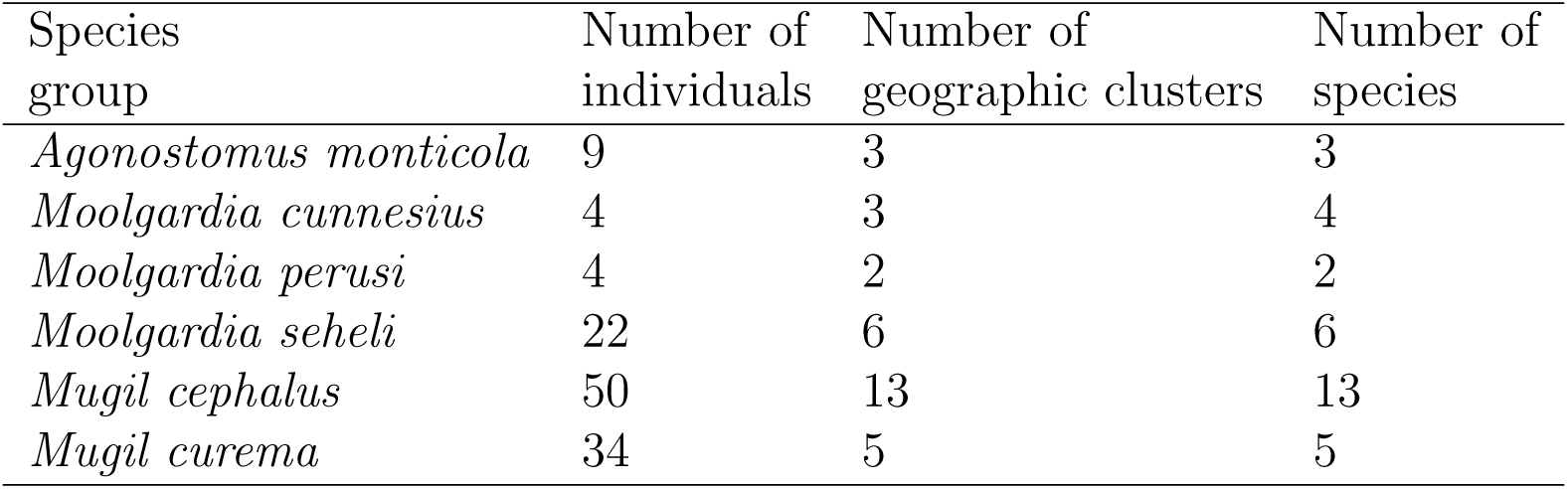
Species delimitation and geographic isolation. Most (92%) of the newly delimited mullet species were identified from geographically isolated species clusters described in previous studies [21, 24].

| Species group | Number of individuals | Number of geographic clusters | Number of species |
| --- | --- | --- | --- |
| *Agonostomus monticola* | 9 | 3 | 3 |
| *Moolgardia cunnesius* | 4 | 3 | 4 |
| *Moolgardia perusi* | 4 | 2 | 2 |
| *Moolgardia seheli* | 22 | 6 | 6 |
| *Mugil cephalus* | 50 | 13 | 13 |
| *Mugil curema* | 34 | 5 | 5 |

### Mullet phylogeny and divergence times

#### Mullet phylogeny

Our analysis recovered two main mullet lineages. The first clade (Subclade A) includes all species currently assigned to the genus *Mugil*, which is paraphyletic with respect to *Xenomugil thoburni*. The second, larger clade (Subclade B) includes all remaining mullets. This result appears quite robust, as we recovered these two subclades in analyses of all three datasets (comprising 233, 100, and 64 sequences) under all of the substitution and relaxed-clock models that we explored (Figures 1, S1, S4, S6, S7). The degree and pattern of uncertainty differ in the two mullet subclades. In Subclade A, all but one of the deeper nodes are strongly supported (*i.e.*, with posterior probability ≥ 0.95), but shallower nodes within the *Mugil cephalus* and *Mugil curema* species complexes are generally uncertain. The situation is reversed in Subclade B, where deeper nodes—particularly along the ‘backbone’ of this subclade—are generally poorly supported, but more recent divergences are generally strongly supported (Figures S4, S6). Despite the considerable phylogenetic uncertainty within Subclade B, the trees inferred from the 100- and 64-species datasets are largely concordant: 53% of the internal nodes occur in both summary trees.

Our results largely accord well with those of previous studies. These two mullet subclades were previously identified in both the large-scale phylogenetic studies of teleost [17] and vertebrate [18] divergence times. In the teleost study—which included 10 mullets among the 1400 species sequenced for 20 nuclear genes and a single mitochondrial gene—the genus *Mugil* was inferred to be the sister group of the remaining mullet species with the exception of *Neomyxus leuciscus* (which was inferred to be sister to *Mugil* and all other mullets). Similarly, the vertebrate study—which included 24 mullets—again identified *Mugil* as the sister group of the remaining mullet species with the exception of *Agonostomus tefairini* (which was inferred to be sister to *Mugil* and all other mullets).

By contrast, our results differ somewhat from those of previous studies based on mitochondrial sequence data. Rather than two major mullet subclades, the Durand *et al.* study [21] identified a number of relatively depauperate lineages (*Sycamugil* + *Rhinomugil*, *Trachystoma*) that formed sequential sister lineages to the remaining mullet species. In that phylogeny, the genus *Mugil* was nested within our Subclade B as sister to a subclade comprising *Agonostomus calatai*, *Joturus pichardi* and *Agonostomus monticola*. In agreement with that study [21], however, our results indicate both that the majority of conventionally recognized mullet genera are not monophyletic, and also that relationships among many lineages within Subclade B remain poorly resolved.

#### Mullet divergence times

We inferred mullet divergence times using two datasets—based on the sample of formally delimited and conventionally recognized mullet species (the 100- and 64-species datasets, respectively)—and analyzed both datasets under six relaxed-clock models—comprising all combinations of the three branch-rate models (RLC, UCLN, UCEX), the two partitioned site models (PS1, PS2), and the single node-age model (SBD). Here we explore the impact of these various species-sampling schemes and relaxed-clock models on estimated divergence times by focussing on the inferred ages of four key nodes: (1) the mullet stem age; (2) the mullet crown age; (3) the crown age of Subclade A, and; (4) the crown age of Subclade B (Tables 8, S6).

For a given relaxed-clock model, the inferred ages of the four key nodes are on average 44% older for the 100-species versus the 64-species dataset. We note, however, that this effect is largely driven by the disparity in divergence-time estimates under the RLC branch-rate model. However, we remain somewhat skeptical of these divergence-time estimates, as our MCMC simulations under the RLC model tended to mix poorly (which is common for this branch-rate model [39]). When the RLC branch-rate model is excluded, the inferred ages of the four key nodes are on average only ∼ 4% older for the 100-species versus the 64-species dataset.

Species sampling imparts both direct and indirect effects on divergence-time estimates. Increased species sampling directly impacts divergence-time estimates by reducing the ‘node-density effect’ [75–77]. This effect causes the lengths of long branches to be disproportionately underestimated; increasing the density of species sampling reduces this bias by breaking up long branches. Because terminal branches are anchored in the present, the increased branch-length estimates conferred by increased species sampling effectively results in older estimates for the ages of internal nodes. Moreover, species sampling *indirectly* impacts divergence-time estimates by influencing the choice of relaxed-clock model. Altering the sample of included species may change the pattern and magnitude of substitution-rate variation across branches, and these different patterns may be best described by different branch-rate models. In fact, different branch-rate models were selected for the two mullet datasets: the UCEX branch-rate model was preferred for the 100-species dataset, whereas the UCLN branch-rate model was preferred for the 64-species dataset (Tables 4, S5). As we discuss below, the choice of branch-rate model may strongly impact of divergence-time estimates.

We observed a strong impact of relaxed-clock models on our estimates of mullet divergence times, and the components of the relaxed-clock models differed in their relative influence on divergence-time estimates. The choice of partition scheme had a pronounced impact divergence-time estimates: ages of the four key nodes inferred under alternative partition schemes (for a given branch-rate model) differed on average by 8.3% and 7.5% for the 100- and 64-species datasets, respectively (Tables 8, S6). The choice of branch-rate model had the most extreme impact divergence-time estimates: ages of the four key nodes inferred under alternative branch-rate models (for a given partition scheme) differed on average by 24.7% and 27.3% for the 100- and 64-species datasets, respectively.

Branch-rate models differ in their ability to capture local fluctuations in substitution-rate variation across adjacent branches. The RLC branch-rate model assumes that substitution rates are locally constant, the UCLN model assumes that rates on adjacent branches are independent and identically distributed (iid) samples from a shared lognormal distribution, and the UCEX model assumes that rates on adjacent branches are iid samples from a shared exponential distribution. Accordingly, extreme fluctuations in substitution rate across branches are best captured by the UCEX > UCLN > RLC branch-rate models [39]. The degree of substitution-rate variation in the 100-species dataset therefore appears to be more pronounced than that in the 64-species dataset, as evidenced by the decisive preference for the UCEX branch-rate model in the former and the UCLN model in the latter. Interaction between species sampling and branch-rate models leads to an apparent paradox. Although mullet divergence times inferred under a *given* relaxed-clock model are on average *older* for the 100-species dataset, the inferred ages for the 100-species dataset are nevertheless *younger* than those for the 64-species dataset under the *preferred* relaxed-clock models (UCEX and UCLN, respectively; Figures 1, S7, Tables 8, S6).

Previous studies have inferred divergence times for two nodes that are recovered in our study—the mullet crown and stem nodes—which provide a basis for comparison. We inferred that mullets diverged from their acanthomorph relatives either 64 Ma (95% HPD [44,90] Ma; Figure 1, Table 8) or 77 Ma (95% HPD [54,107] Ma; Figure S7, Table S6) based on analyses of the 100- or 64-species datasets, respectively. A Late Cretaceous/Early Paleogene origin for mullets is generally consistent with recent large-scale studies of teleost divergence times. For example, Near *et al.* [18] inferred a similar stem age for mullets (∼77 Ma), whereas Betancur *et al.* [17] inferred a somewhat older mullet stem age (∼89 Ma). We inferred the earliest divergence within mullets—which gave rise to the two main subclades—occurred either 55 Ma (95% HPD [41,72] Ma) or 65 Ma (95% HPD [50,83] Ma) based on analyses of the 100- or 64-species datasets, respectively. Our estimate of the crown age of mullets is similar to that based on a supermatrix analysis of vertebrates (∼60 Ma) [19], but is somewhat older than estimates of the study by McMahan *et al.* [25] (41.5 Ma), and Betancur *et al.* [17] (44.5 Ma). We suspect that these discrepancies stem from disparities in taxon sampling and fossil calibration: our analysis included 98 (or 62) mullet species and three fossil calibrations, whereas McMahan *et al.* [25] included seven mullet species and a single fossil calibration, and Betancur-R *et al.* [17] included 10 mullet species and no mullet fossil calibrations.

**Table 8.**
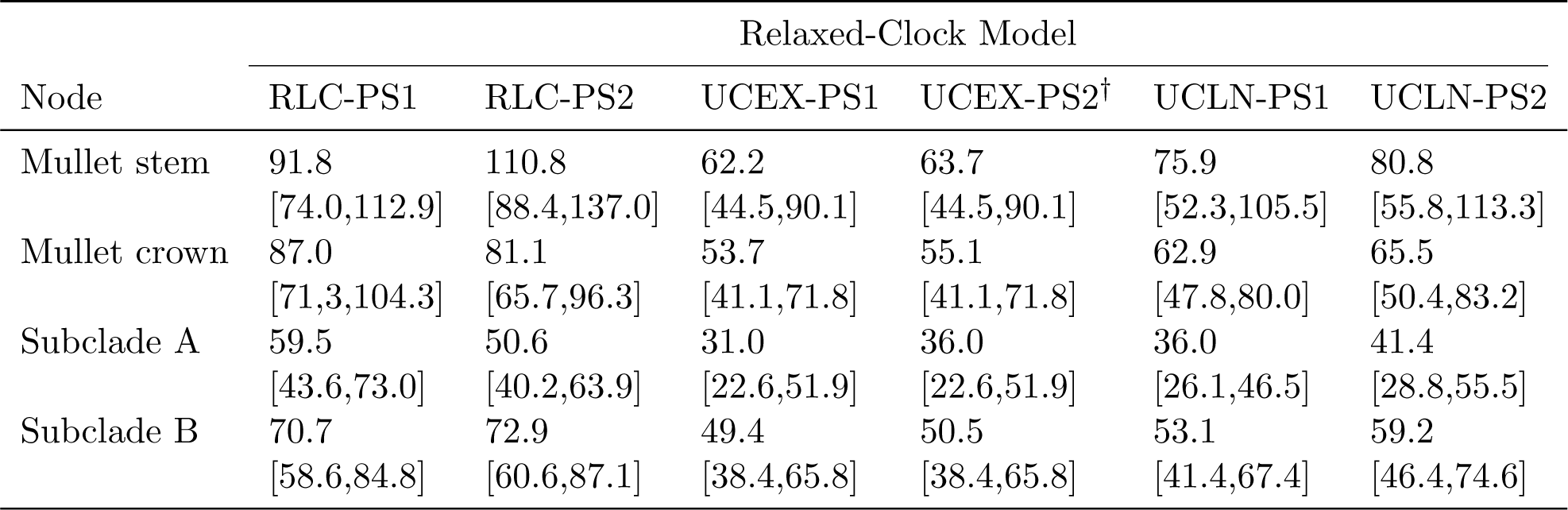
The impact of relaxed-clock models on divergence-time estimates. Comparisons are based on analyses of the 100-species dataset. We report the estimated median [and 95% HPD] of ages for four key nodes under the six relaxed-clock models that we explored. ^†^The divergence-time estimates under the preferred relaxed-clock model.

| Node | Relaxed-Clock Model |  |  |  |  |  |
| --- | --- | --- | --- | --- | --- | --- |
|  | RLC-PS1 | RLC-PS2 | UCEX-PS1 | UCEX-PS2<sup>†</sup> | UCLN-PS1 | UCLN-PS2 |
| Mullet stem | 91.8<br>[74.0,112.9] | 110.8<br>[88.4,137.0] | 62.2<br>[44.5,90.1] | 63.7<br>[44.5,90.1] | 75.9<br>[52.3,105.5] | 80.8<br>[55.8,113.3] |
| Mullet crown | 87.0<br>[71.3,104.3] | 81.1<br>[65.7,96.3] | 53.7<br>[41.1,71.8] | 55.1<br>[41.1,71.8] | 62.9<br>[47.8,80.0] | 65.5<br>[50.4,83.2] |
| Subclade A | 59.5<br>[43.6,73.0] | 50.6<br>[40.2,63.9] | 31.0<br>[22.6,51.9] | 36.0<br>[22.6,51.9] | 36.0<br>[26.1,46.5] | 41.4<br>[28.8,55.5] |
| Subclade B | 70.7<br>[58.6,84.8] | 72.9<br>[60.6,87.1] | 49.4<br>[38.4,65.8] | 50.5<br>[38.4,65.8] | 53.1<br>[41.4,67.4] | 59.2<br>[46.4,74.6] |

### Evolution of feeding preference in mullets

#### Evolution of diet in mullets

We inferred the evolution of diet—as a discrete trait with three states (algae, detritus, or invertebrates)—using the trees inferred from both the 100-species and 64-species datasets. For each tree, we inferred diet evolution under six discrete-trait models. These models comprised all possible combinations of two continuous-time Markov models (that describe the relative rates of change among the three discrete diet states) and three branch-rate models (that describe how rates of diet evolution vary across branches of the phylogeny). Here, we discuss implications of the preferred discrete-trait models for the evolution of feeding preference in mullets.

Both the 100- and 64-species datasets decisively preferred the CTM 6-rate model (Tables 6, S7), which accommodates asymmetric instantaneous rates of change between each pair of states (*i.e.*, *q*_*ij*_ ≠ *q*_*ji*_). This implies that, for both trees, pairwise rates of change between diets are unequal. In fact, this is clear from the estimated instantaneous rates; the absolute difference between the forward and reverse instantaneous rates of change between states—that is, |*q*_*ij*_ − *q*_*ji*_ | for each pair of diets *i* and *j*—was inferred to differ by an average of 39.7% and 32.8% in the 100- and the 64-species datasets, respectively (Tables 9, S8).

In contrast to the continuous-time Markov component of the discrete-trait models—where both datasets preferred the same asymmetric CTM 6-rate model—the 100- and 64-species datasets preferred different branch-rate models. Specifically, rates of diet evolution across branches of the the 64-species tree were best described by the constant-rate morphological clock (CRMC) branch-rate model (Table S7). This implies that rates of substitution and diet evolution vary independently across branches of the 64-species tree: rates of substitution vary across lineages under the UCLN branch-rate model, whereas rates of diet evolution are constant through time and across lineages of the tree. This situation contrasts sharply with that of the 100-species tree, where rates of substitution and diet evolution covary across branches under a shared UCEX branch-rate model (Table 6).

Presumably, the preference for the shared branch-rate model by the 100-species dataset stems from a key aspect of the species sampling. Specifically, the 100-species dataset adds several clusters of newly delimited species that tend to be characterized by low evolutionary rates. Consider, for example, the clade of 13 distinct species of *Mugil cephalus*. Each species exhibits the same algal diet (implying that the rate of diet evolution within this clade is very low), and the pairwise sequence divergence between these newly delimited species is also relatively small (implying that substitution rates within this clade are also low). Accordingly, rates of substitution and diet evolution tend to be low within the clusters of newly delimited species, which drives the preference for a shared branch-rate model in the 100-species dataset.

Ascertaining whether rates of phenotypic and molecular evolution covary across branches is pertinent to recently proposed ‘tip-dating’ methods [78, 79]. This approach for estimating divergence times accommodates uncertainty in the placement of fossil calibrations by jointly estimating the phylogenetic position of fossils (and the duration of branches connecting them to the tree) from datasets comprising partitions of both nucleotide sequences and discrete morphological traits. Critically, this ‘tip-dating’ approach assumes that rates of substitution and morphological evolution covary under a shared branch-rate model. Our results suggest that this assumption may not always be valid.

Despite obvious differences in the two datasets—and consequent differences in the preferred trait model—some aspects of the inferred history of diet evolution are similar for the 100- and 64-species datasets. For both datasets, invertebrates were inferred to comprise the ancestral diet (although with slightly more uncertainty in the 100-species dataset), which gave rise to multiple independent origins of the algal and detritivorous diets (Figures 2, S8). Our estimates of the overall and relative number of changes between diets—inferred using the Markov-jump approach [66, 67]—differed for the the two mullet datasets: the total count of diet changes was somewhat higher for the 100- versus the 64-species dataset (35.7 vs. 26.4 changes, respectively; Tables 9, S8). [We note that the higher inferred rates of diet evolution contribute to the increased uncertainty in ancestral states in the 100-species dataset.] Average counts of changes between diets inferred for the 100- and 64-species datasets were quite different for transitions between detritus-algae (6.4:0.5), invertebrates-algae (4.5:8.4), and detritus-invertebrates (8.0:0.3). Interestingly—and at first glance, perhaps confusingly—estimates for the average number of complementary changes in diet were quite similar for the 100- and 64-species datasets: algae-detritus (3.0:2.3), algae-invertebrates (1.6:1.2), and invertebrates-detritus (12.2:13.7).

Although seemingly paradoxical, the disparity in the counts of forward and reverse transitions between complementary diets in the 100- and 64-species datasets stems from a quirk of species sampling in our study. As described previously, the 100-species dataset adds several clusters of newly delimited species that exhibit identical diets; this impacts estimates of the instantaneous rates and counts of changes between diets.

**Table 9.** Inferred rates and counts of dietary change in mullets. We inferred the evolution of diet in mullets under the preferred discrete-trait model (Table 6) with six instantaneous-rate parameters, *q*_*ij*_, between the three states—algae (A), detritus (D), and invertebrates (I)—and estimated the expected number of changes between states using the Markov-jump approach.

| Instantaneous-rate parameter | Mean rate | 95% HPD rate | Mean count | 95% HPD count |
| --- | --- | --- | --- | --- |
| $q_{AD}$ | 0.81 | [7.30E-3, 1.98] | 3.00 | [0.69, 6.31] |
| $q_{DA}$ | 0.53 | [3.15E-5, 1.45] | 6.37 | [1.42, 11.99] |
| $q_{AI}$ | 1.28 | [1.80E-3, 2.91] | 1.59 | [2.71E-5, 3.97] |
| $q_{IA}$ | 1.17 | [7.67E-2, 2.63] | 4.53 | [1.06E-3, 9.21] |
| $q_{ID}$ | 0.64 | [6.15E-4, 1.59] | 12.18 | [4.56, 22.16] |
| $q_{DI}$ | 1.48 | [1.89E-1, 3.19] | 8.00 | [1.14E-5, 15.89] |

#### Correlates of diet evolution in mullets

We explored evolutionary correlations between feeding preference and several other variables in mullets under the latent-liability model [69]. Specifically, we evaluated all pairwise correlations between two continuous traits—trophic index and total length—and two discrete traits—habitat type (marine, non-marine), and geographic distribution (tropical, non-tropical). We performed these analyses both for the 100- and 64-species datasets, and for both datasets we repeated the analyses over a range of (hyper)prior values to assess the robustness of any inferred correlations.

Our analyses of the 100-species dataset revealed three significant correlations (Figure 3): (1) a positive correlation between trophic index and habitat (with herbivorous and/or detritivorous species predominantly occurring in marine habitats); (2) a negative correlation between trophic index and geographic distribution (with herbivorous species occurring predominantly in subtropical and temperate regions), and; (3) a negative correlation between body size and geographic distribution (with larger species occurring predominantly in subtropical and temperate regions). Our sensitivity analyses indicate that these results are robust to all nine combinations of the (hyper)prior values that we explored (Figure S5). Taken at face value, the inferred correlations for the 100-species dataset suggest that the opportunities for evolution of herbivorous and/or detritivorous diets have been more favorable in relatively large mullet species residing in colder (temperate/subtropical) marine habitats.

**Figure 3.**
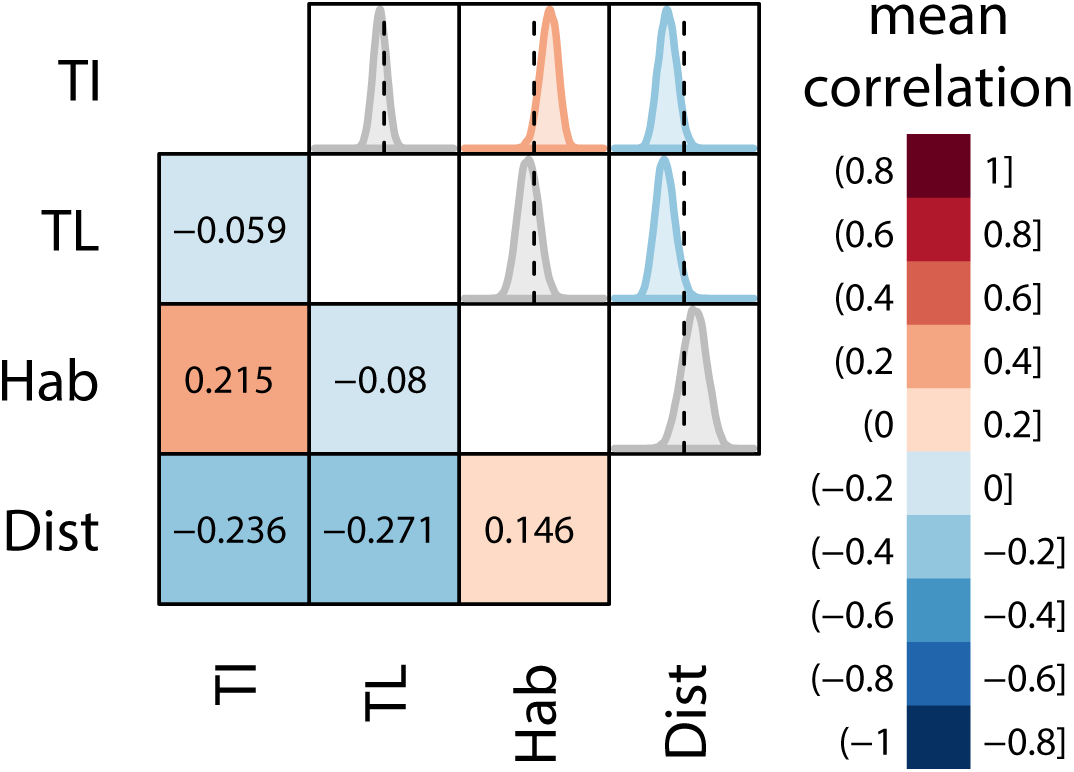
Correlates of diet evolution in mullets. Estimates are based on analyses of the 100-species dataset under the latent-liability model [69]. We explored correlations between two discrete and two continuous traits. These traits (abbreviations) [and states] include: trophic index (TI); total length (TL) [centimeters]; habitat type (Habitat) [marine, non-marine]; and distribution (Distribution) [tropical, non-tropical]. The lower diagonal depcts the mean correlation coefficients for each pair of traits (see inset legend), and the upper diagonal depicts the corresponding marginal densities of the correlation coefficients (values range from −1 to +1). Densities are colored according to their mean value only if they differ significantly from zero (*i.e.*, the posterior probability that the value is equal to or more extreme than 0 is < 0.05).

Our analyses of the 64-species dataset, however, suggest that these findings—like those for divergence times and diet evolution—are somewhat sensitive to the set of distinct species included in the analysis. In our analyses of the 64-species dataset, the correlation between trophic index and habitat becomes marginally non-significant, and that between trophic index and geographic distribution is rendered non-significant. Our failure to recover significant correlations in the 64-species dataset that were identified in the 100-species dataset likely reflects two factors. First, the power to detect correlations under the latent-liability model scales with sample size: the method is therefore more likely to detect correlations in larger trees. Moreover, the 100-species dataset—as noted previously—adds several clusters of newly delimited species that are phenotypically identical. Within these clusters of identical species, all of the variables will necessarily be perfectly correlated, which may lead to the identification of significant correlations across the entire tree. Assuming that we have correctly delimited species, this effect simply increases our power to detect bonafide correlations between traits. Conversely, if our species delimitation is invalid (particularly if species have been overly ‘split’), then this effect will induce a bias causing the identification of spurious correlations between traits.

Of the correlations detected in the 100-species dataset, only that between body size and geographic distribution remains significant for the 64-species dataset (Figure S9). Accordingly, the positive correlation between body size latitude detected in mullets is consistent with Bergmann’s rule [80]. Although originally proposed for endothermic organisms, this classic (and controversial) ecogeographic principle has also been reported in ectotherms [81], including several groups of freshwater and marine fishes [82–87].

### Conclusions

We identified 98 distinct mullet species within the 233-sequence dataset sampled from 62 nominal species. Most of the newly delimited species correspond to geographically isolated lineages, suggesting these species arose by (or are currently undergoing) the process of allopatric speciation. We performed a parallel series of comprehensive statistical analyses on the formally delimited (100-species) and conventionally recognized (64-species) datasets to estimate phylogenetic relationships, divergence times, and the evolution of feeding preference in mullets.

Several results appear robust to the choice of species sample: mullets diverged from other acanthomorphs in the Late Cretaceous/Early Paleogene and today are distributed among two clades—the first mainly comprised of *Mugil* species, the second containing the remaining species. Moreover, it is clear that the characteristic diet of mullets—on algae and detritus—arose independently multiple times from an ancestral diet on invertebrates. Similarly, it appears that body size in mullets increases with latitude, as would be predicted by Bergmann’s rule.

By contrast, many of our findings are somewhat sensitive to the set of recognized mullet species. For example, almost half of the shared internal nodes differ in the trees inferred from the 100- and 64-species datasets, and—for a given relaxed-clock model—divergence-time estimates for the 100-species dataset are somewhat older than those for the 64-species dataset. Similarly, the relative rates and average counts of changes between diets differ substantially between the 100- and 64-species datasets. Moreover, two of the three identified correlations between traits—that herbivorous and/or detritivorous species predominantly occur in marine habitats, and that herbivorous species predominantly occur in subtropical and temperate regions—depend critically on the definition of mullet species adopted for these analyses.

Our study also emphasizes the critical impact of model choice in statistical phylogenetic analyses. The choice of relaxed-clock model, for example, had an even more dramatic impact on divergence-time estimates than did the sample of mullet species included in these analyses. The results of our mullet study—albeit anecdotal—highlight the importance of carefully evaluating and rigorously selecting among candidate relaxed-clock model in studies of species divergence times.

Although we have explored the sensitivity of statistical phylogenetic inferences—on estimates of phylogeny, divergence times and trait evolution—to *species delimitation* in mullets, we suspect that our findings are also relevant to the more general issue of incomplete and/or non-random *species sampling* in comparative studies for groups where species boundaries are uncontroversial.

## Acknowledgments

We are grateful to Gabriela Cybis for providing access to code implementing the latent-liability model, to Andrew Magee for assistance with analyses, to Mark Miller for providing access to computational resources via CIPRES, and to Peter Wainwright for thoughtful discussion. This research was supported by NSF grants DEB-0842181, DEB-0919529, and DBI-1356737 awarded to BRM. Computational resources for this work were provided by an NSF XSEDE grant (DEB-120031) to BRM.

## Supporting Information

### Data/model files archived on the Dryad Digital Repository

#### Data Availability Statement

The authors confirm that all data supporting the results of this study are fully available without restriction. We have made provided all of the molecular and comparative data as input files (with the corresponding full model specifications) that were used to perform the analyses described in our study. These files have been deposited to the Dryad database. The Dryad data identifier for this study is: doi:10.5061/dryad.h26v3.

1. The 233-sequence alignment in NEXUS format with `MrBayes` model block used to infer the phylogram depicted in Figure S1 (`mullet 233.nex`).
2. The 100-species alignment in NEXUS format with `MrBayes` model block used to infer the phylogram depicted in Figure S4 (`mullet 100.nex`).
3. The 100-species alignment in XML format with the selected relaxed-clock model used to infer the dated phylogeny depicted in Figure 1 (`mullet 100 AIC UCEX.xml`).
4. The 100-species alignment in XML format with the selected discret-trait model used to infer the history of diet evolution depicted in Figure 2 (`mullet 100 UCEXs CTM6.xml`).
5. The 100-species trait dataset in tab-delimited text format file used to infer the the history of diet evolution depicted in Figure 2 (`mullet 100 diet traits.txt`).
6. The 100-species alignment in XML format with the selected latent-liability model used to infer character correlation depicted in Figures 3 and S5 (`mullet 100 latent.xml`).
7. The 64-species alignment in NEXUS format with MrBayes model block used to infer the phylogram depicted in Figure S6 (`mullet 64.nex`).
8. The 64-species alignment in XML format with the selected relaxed-clock model used to infer the dated phylogeny depicted in Figure S7 (`mullet 64 AIC UCLN.xml`).
9. The 64-species alignment in XML format with the selected discret-trait model used to infer the history of diet evolution depicted in Figure S8 (`mullet 64 CRMC CTM6.xml`).
10. The 64-species trait dataset in tab-delimited text format file used to infer the the history of diet evolution depicted in Figure S8 (`mullet 64 diet traits.txt`).
11. The 64-species alignment in XML format with the selected latent-liability model used to infer character correlation depicted in Figure S9 (`mullet 64 latent.xml`).
12. A report diagnosing the MCMC performance of analyses of the 100-species dataset under the preferred relaxed-clock model as a PDF file (`mullet 100 MCMC.pdf`).
13. A report diagnosing the MCMC performance of the 64-species dataset under the preferred relaxed-clock model as a PDF file (`mullet 64 MCMC.pdf`).

## Analyses of the 233-sequence dataset

In order to estimate the number of distinct mullet species, we estimated a rooted phylogram from all available sequence data for the three mitochondrial gene regions. We first selected a mixed model that provided the best fit to 8 predefined data subsets using `PartitionFinder` [31] (Table S1).

**Table S1.** Mixed-model selection for the 233-sequence dataset. We selected among the space of partition schemes that variously assign substitution models to 8 data subsets using both the BIC and AIC model-selection methods implemented in `PartitionFinder`. Substitution models that are linked across multiple data subsets are indicated with superscripts. The number of free substitution-model parameters (excluding branch lengths) for each of the partition schemes is indicated in the rightmost column.

| Scheme | Data Subset |  |  |  |  |  |  |  |  |
| --- | --- | --- | --- | --- | --- | --- | --- | --- | --- |
|  | *cox1* |  |  | *cytb* |  |  | 16S |  | NP |
|  | 1<sup>st</sup> pos. | 2<sup>nd</sup> pos. | 3<sup>rd</sup> pos. | 1<sup>st</sup> pos. | 2<sup>nd</sup> pos. | 3<sup>rd</sup> pos. | stem | loop |  |
| PS1(BIC) | F81+$\Gamma$ | GTR+$\Gamma$ | SYM+$\Gamma^1$ | SYM+$\Gamma$ | HKY+$\Gamma$ | GTR+$\Gamma$ | SYM+$\Gamma^1$ | GTR+$\Gamma$ | 54 |
| PS2(AIC) | GTR+I | GTR+$\Gamma$ | GTR+$\Gamma$ | SYM+$\Gamma$ | GTR+$\Gamma$ | GTR+$\Gamma$ | SYM+$\Gamma$ | GTR+$\Gamma$ | 73 |

We then estimated the posterior probability distribution of trees (and other model parameters) under the selected partition scheme using the MCMC algorithm implemented in `MrBayes` [32]. We summarized the resulting posterior distribution of phylogenies as an all-compatible majority-rule consensus tree, which we rooted using the two surfperch outgroup species (Figure S1). The status of the nominal mullet species in this phylogeny are summarized in Figure S2.

### Nominally monophyletic

Several species were represented by a single accession in the 233-sequence dataset, and so are trivially monophyletic: *Agonostomus catalai*, *Cestraeus goldiei*, *Cestraeus oxyrhyncus*, *Chelon melinopterus*, *Liza abu*, *Liza klunzingeri*, *Liza parsia*, *Liza saliens*, *Liza tricuspidens*, *Mugil gyrans*, *Myxus elongatus*, *Oedalechilus labiosus*, *Paramugil parmatus*, *Rhinomugil corsula*, *Sicamugil cascasia*,*Sicamugil hamiltonii*, *Trachystoma petardi*, and *Valamugil speigleri*.

### Monophyletic

The following 35 species were inferred to be monophyletic: *Agonostomus monticola*, *Aldrichetta forsteri*, *Chaenomugil proboscideus*, *Chelon labrosus*, *Crenimugil crenilabis*, *Ellochelon vaigiensis*, *Joturus pichardi*, *Liza affinis*, *Liza argentea*, *Liza aurata*, *Liza bandialensis*, *Liza carinata*, *Liza dumerili*, *Liza falcipinnis*, *Liza grandisquamis*, *Liza haematocheila*, *Liza ramado*, *Liza richardsonii*, *Moolgarda engeli*, *Moolgarda perusii*, *Mugil bananensis*, *Mugil capurrii*, *Mugil chelo*, *Mugil hospes*, *Mugil incilis*, *Mugil liza*, *Mugil rubrioculus*, *Mugil trichodon*, *Myxus capensis*, *Neomyxus leuciscus*, *Oedalechilus labeo*, *Rhinomugil nasutus*, *Valamugil buchanani*, *Valamugil robustus*, and *Xenomugil thoburni*.

### Paraphyletic

Six species were inferred to be paraphyletic: *Chelon labrosus* (with respect to *Mugil chelo*), *Chelon macrolepis* (with respect to *Liza carinata*), *Moolgarda cunnesius* (with respect to *Valamugil speigleri*), *Moolgarda seheli*, (with respect to *Crenimugil crenilabis*), *Mugil cephalus* (with respect to *Mugil liza*), and *Mugil curema* (with respect to *Mugil gyrans* and *Xenomugil thoburni*). *Polyhyletic*—Only two species were inferred to be polyphyletic: *Chelon planiceps*, and *Liza alata*.

**Figure S1.**
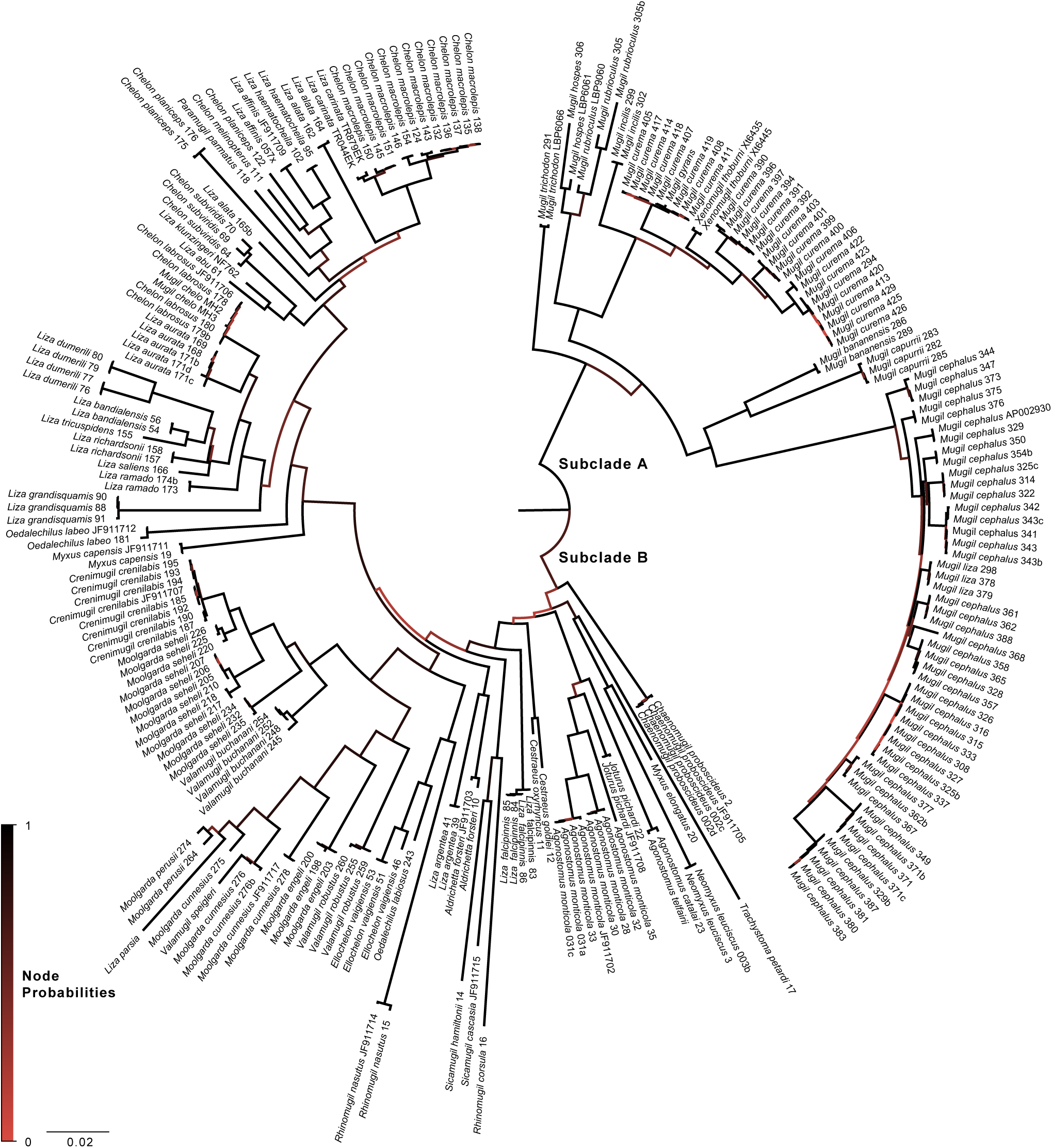
Bayesian estimate of mullet phylogeny for the 233-sequence dataset. Branch lengths are rendered proportional to the expected number of substitutions per site (see inset scale). The color or internal branches reflects the posterior probability of the corresponding nodes (see inset legend). Two major subclades are inferred: Subclade A is primarily comprised of *Mugil* species; Subclade B includes the remaining mullet species.

**Figure S2.**
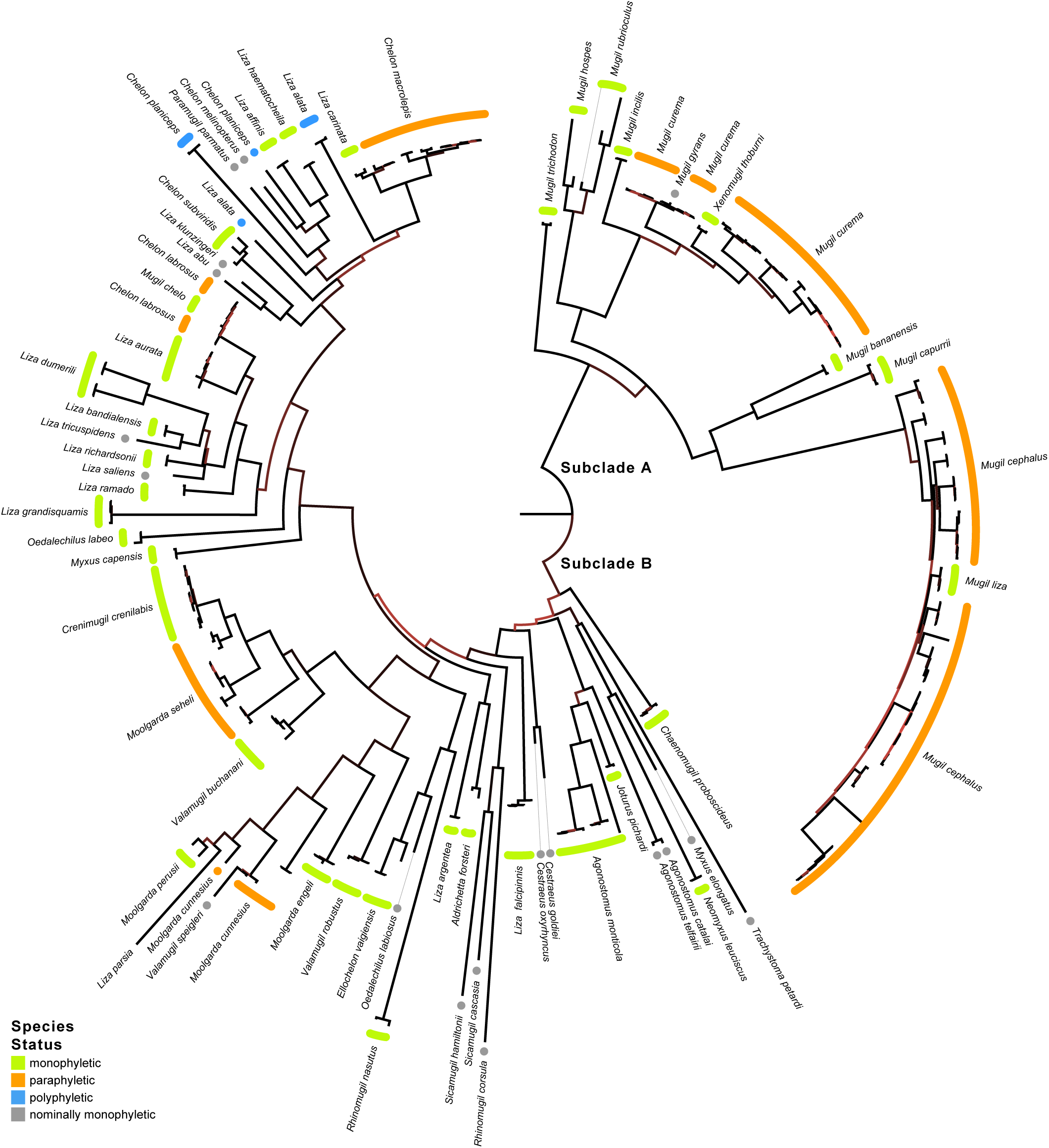
The implied status of mullet species in the 233-sequence phylogeny. The Bayesian estimate of phylogeny for the 233-sequence dataset (Figure S1) emphasizing the status of nominal mullet species (see inset legend).

**Figure S3.**
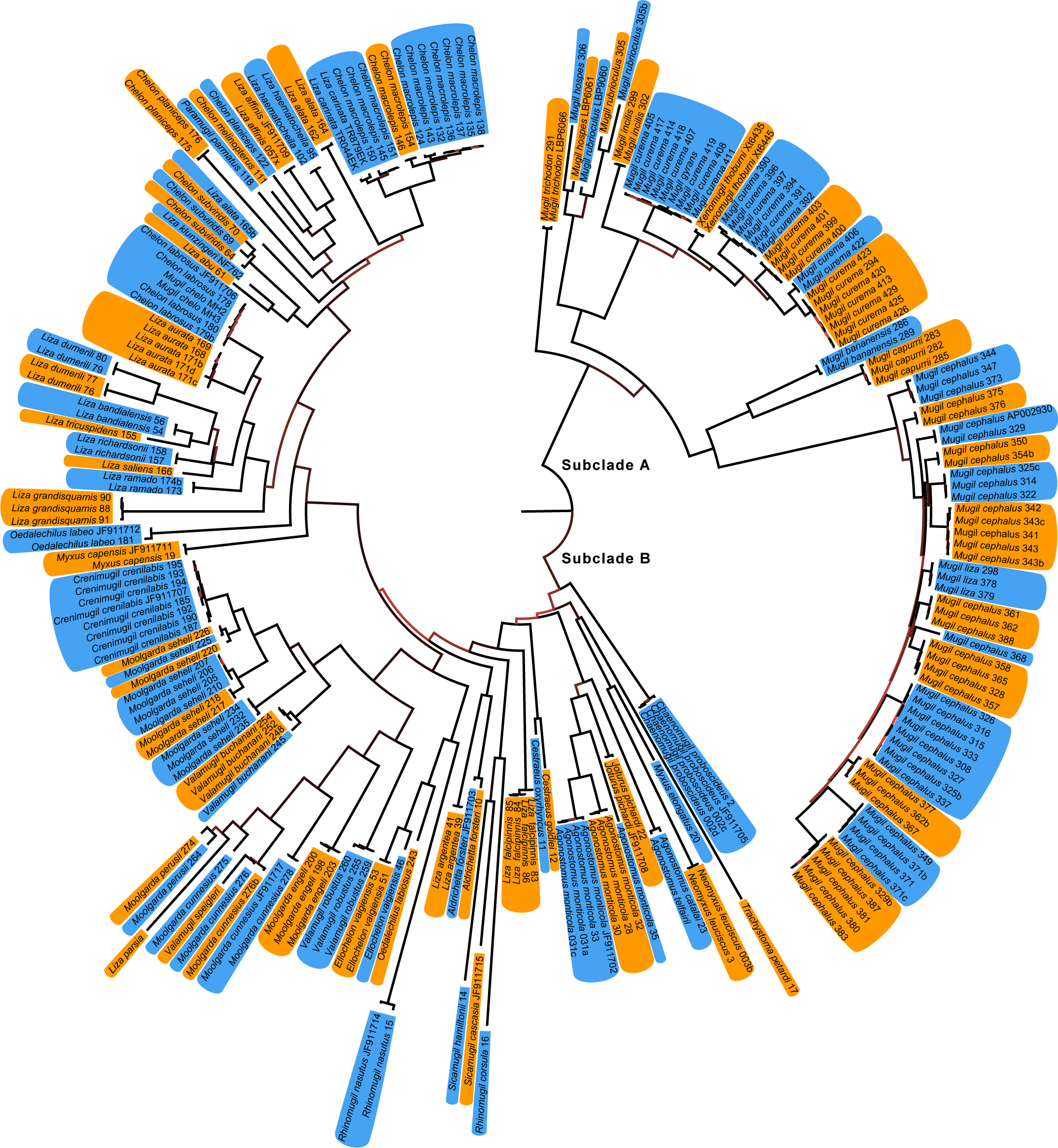
Delimitation of mullet species in the 233-sequence phylogeny. The 98 distinct mullet species delimited from the 233-sequence dataset (Figure S1) using the Poisson tree process model are indicated in alternating blue and orange colors.

## Analyses of the 100-species dataset

**Figure S4.**
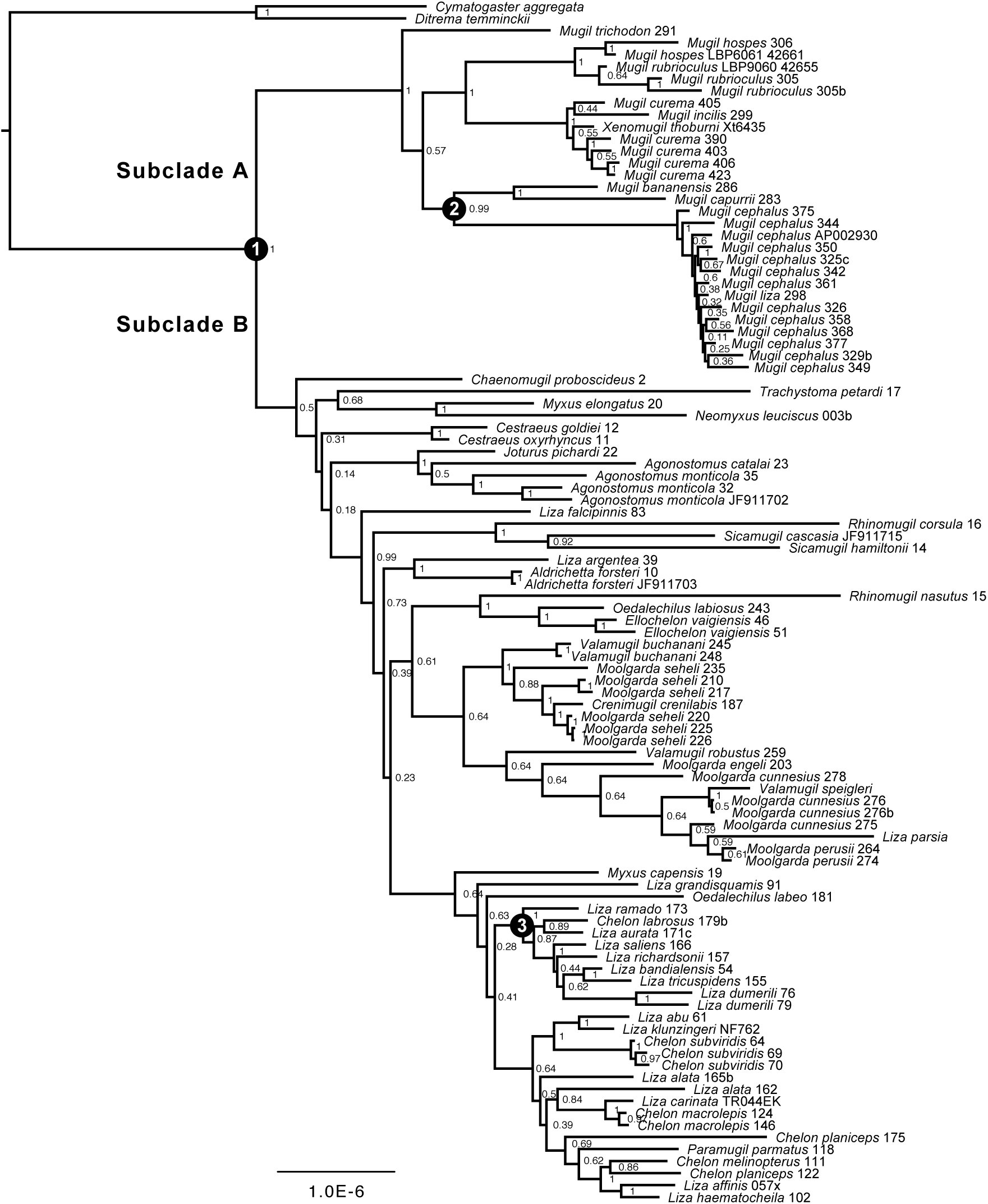
Bayesian estimate of mullet phylogeny. Estimates are based on the 100-species dataset under the preferred partition scheme (PS1; Table 1). Branch lengths are rendered proportional to the expected number of substitutions per site (inset scale bar). Numbers adjacent to internal nodes indicate the corresponding marginal probabilities; the three circled nodes indicate the location of the corresponding fossil calibrations (see Table 2).

## Correlated-trait evolution: assessing prior sensitivity

**Figure S5.**
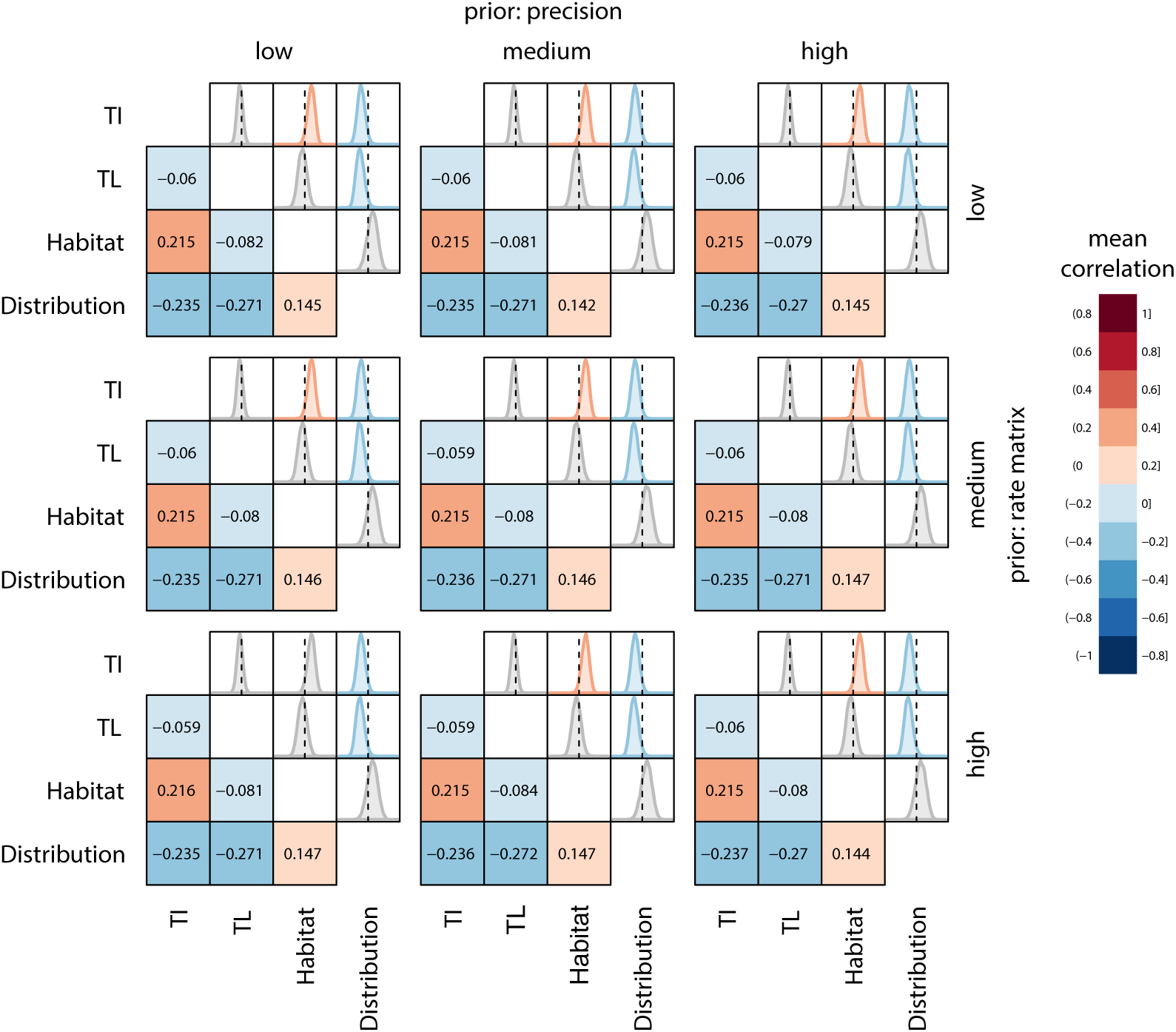
Correlates of diet evolution in mullets. Estimates are based on analyses of the 100-species dataset under the latent-liability model [69]. Traits (abbreviations) [and states] include: trophic index (TI); total length (TL) [centimeters]; habitat type (Habitat) [marine, non-marine]; and distribution (Distribution) [tropical, non-tropical]. We repeated these analyses using nine distinct combinations of priors on the precision-matrix and rate-matrix parameters. Each row of panels corresponds to low, medium, or high values for the rate-matrix parameter; each column corresponds to low, medium and high values for the precision-matrix parameter. Within each panel, the lower diagonal depcts the mean correlation coefficients for each pair of traits (see inset legend), and the upper diagonal depicts the corresponding marginal densities of the correlation coefficients (values range from −1 to +1). Densities are colored according to their mean value only if they differ significantly from zero (*i.e.*, the posterior probability that the value is equal to or more extreme than 0 is < 0.05).

## Analyses of the 64-species dataset

### Sequence data

Despite our efforts, there remains considerable uncertainty regarding the actual number of mullet species. This naturally raises concerns regarding the sensitivity of our findings—regarding the phylogeny, divergence times, and evolution of feeding in mullets—to this critical source of uncertainty. To address this issue, we performed replicate analyses using a dataset for the conventionally recognized mullet species. We defined this dataset by randomly selecting a single sequence for each of the 62 mullet species represented in the 233-sequence dataset. That is, for every species with *N* > 1 sequences, we randomly selected a single sequence (where each sequence was selected with a probability of 1/*N*) without reference to the phylogenetic position of the sequences or the values of other variables (diet, body length, habitat, or geographic distribution).

We aligned the selected sequences for each gene using `MUSCLE` v.3.8.31 [27], confirmed the reading frame by examining the amino-acid translation in `AliView` v.1.18 [28], and then trimmed the ragged 3*′* and 5*′* ends of each aligned gene. The concatenated alignment comprised a total of 1986 sites—including 604 bp of 16S, 598 bp of *cox1*, and 784 bp of cyt*b*—for a total of 64 s (the 62 mullet species and two surfperch species as outgroups; *Cymatogaster aggregata* and *Ditrema temninckii*), with 9.9% missing data.

### Preliminary analyses

Estimates of absolute divergence times typically assign fossil calibrations to one or more nodes as prior probability densities, where the calibrated node is assumed to be monophyletic. To assess support for the calibrated nodes in the 100-species dataset, we performed a series of preliminary analyses under the non-clock tree model using `MrBayes` v.3.2.2. [32]. Specifically, we first selected mixed-substitution models (‘partition schemes’) for the sequence alignment using `PartitionFinder` v.1.1.1 [31]. We defined 8 data subsets—one for each of the three codon positions in the two protein-coding genes, and one each for the stem and loop regions of the 16S ribosomal gene—and used the heuristic (‘greedy’) algorithm to explore the space of partition schemes for the set of substitution models implemented in `MrBayes`. We then used both the Bayesian information criterion (BIC) [33] and the Akaike Information Criterion (AIC) [40] to select among the candidate partition schemes. The two resulting partition schemes—the first selected using the BIC (‘PS1’), and the second using the AIC (‘PS2’)—are depicted in Table S2.

We then estimated the joint posterior probability distribution for each of the candidate mixed-substitution models (PS1 and PS2) by running four replicate MCMC simulations using `MrBayes` v.3.2.2 [32]. We ran each simulation for 10^8^ cycles, and thinned each chain by sampling every 10,000^th^ state. We assessed reliability of the simulations in the usual way, and combined the stationary samples from the four replicate simulations. We then estimated the relative fit of the data to the two partition schemes using Bayes factors. To this end, we first estimated the marginal likelihood for each mixed-substitution model using the posterior simulation-based analog of the AIC through Markov chain Monte Carlo (AICm) [68]; we performed these estimates using the method-of-moments estimator [63, 64] implemented in `Tracer` v.1.6, where we estimated the standard error (S.E.) using 1000 bootstrap replicates. We then compared the fit of the two partition schemes to the data by calculating the Bayes factor as 2*ln*(*M*_1_ : *M*_2_), where *M*_*i*_ is the marginal likelihood for model *i* (Table S3).

**Table S2.** Mixed-model (partition scheme) selection for the 64-species dataset. We selected among the space of partition schemes that variously assign substitution models implemented in `MrBayes` to the 8 pre-specified data subsets using both the BIC and AIC model-selection methods implemented in `PartitionFinder`. Substitution models that are linked across multiple data subsets are indicated with superscripts. The number of free substitution-model parameters (excluding branch lengths) for each partition scheme is indicated in the rightmost column.

| Partition | Data Subset |  |  |  |  |  |  |  | NP |
| --- | --- | --- | --- | --- | --- | --- | --- | --- | --- |
|  | *cox1* |  |  | *cytb* |  |  | 16S |  |  |
|  | 1<sup>st</sup> pos. | 2<sup>nd</sup> pos. | 3<sup>rd</sup> pos. | 1<sup>st</sup> pos. | 2<sup>nd</sup> pos. | 3<sup>rd</sup> pos. | stem | loop |  |
| PS1(BIC) | HKY+I<sup>1</sup> | HKY+Γ | GTR+Γ | SYM+Γ<sup>2</sup> | HKY+I<sup>1</sup> | GTR+Γ | SYM+Γ<sup>2</sup> | SYM+Γ | 49 |
| PS2(AIC) | HKY+I | GTR+Γ | GTR+Γ | SYM+Γ | HKY+Γ | GTR+Γ | K2P+Γ | GTR+Γ | 73 |

**Table S3.**
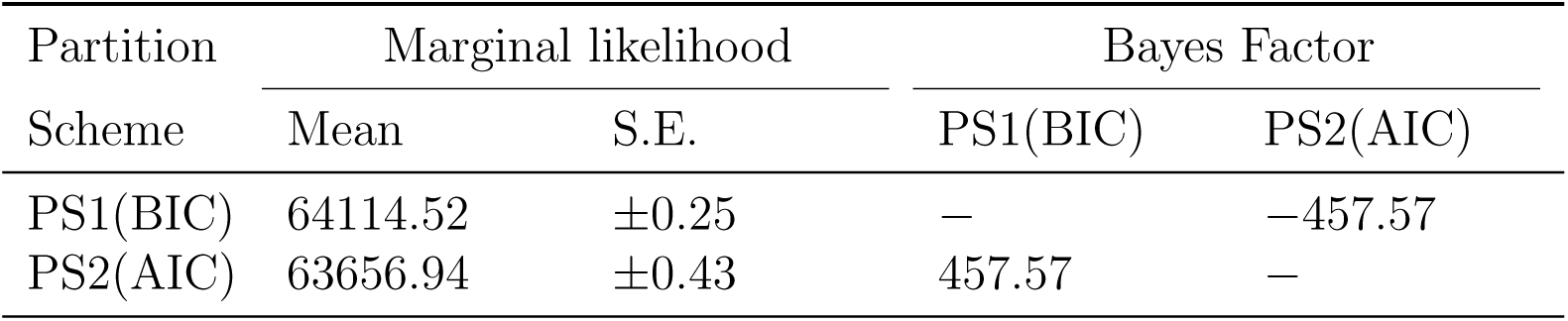
Marginal likelihoods and Bayes factor comparisons for partition schemes. Model comparisons are based on analyses of the 64-species dataset. Marginal likelihoods for each of the candidate partition schemes are based on the AICm method-of-moments estimator [63, 64]. Estimates of the standard error (S.E.) are based on 1000 bootstrap replicates performed in `Tracer` v.1.6. We compared the fit of the two partition schemes to the data using Bayes factors, which we calculated as 2*ln*(*M*_1_ *− M*_2_), where *M*_*i*_ is the marginal-likelihood estimate for partition scheme *i*. The table compares marginal likelihoods for the pair of models in row *i* and column *j*: positive values indicate support for the corresponding model in row *i*. The PS2(AIC) partition scheme is decisively preferred over the PS1(BIC) mixed model (*ln*BF > 4.6) [65].

We summarized the composite marginal posterior probability distribution of trees as an all-compatible majority rule consensus tree (Figure S6), which indicates strong support for the calibration points.

**Figure S6.**
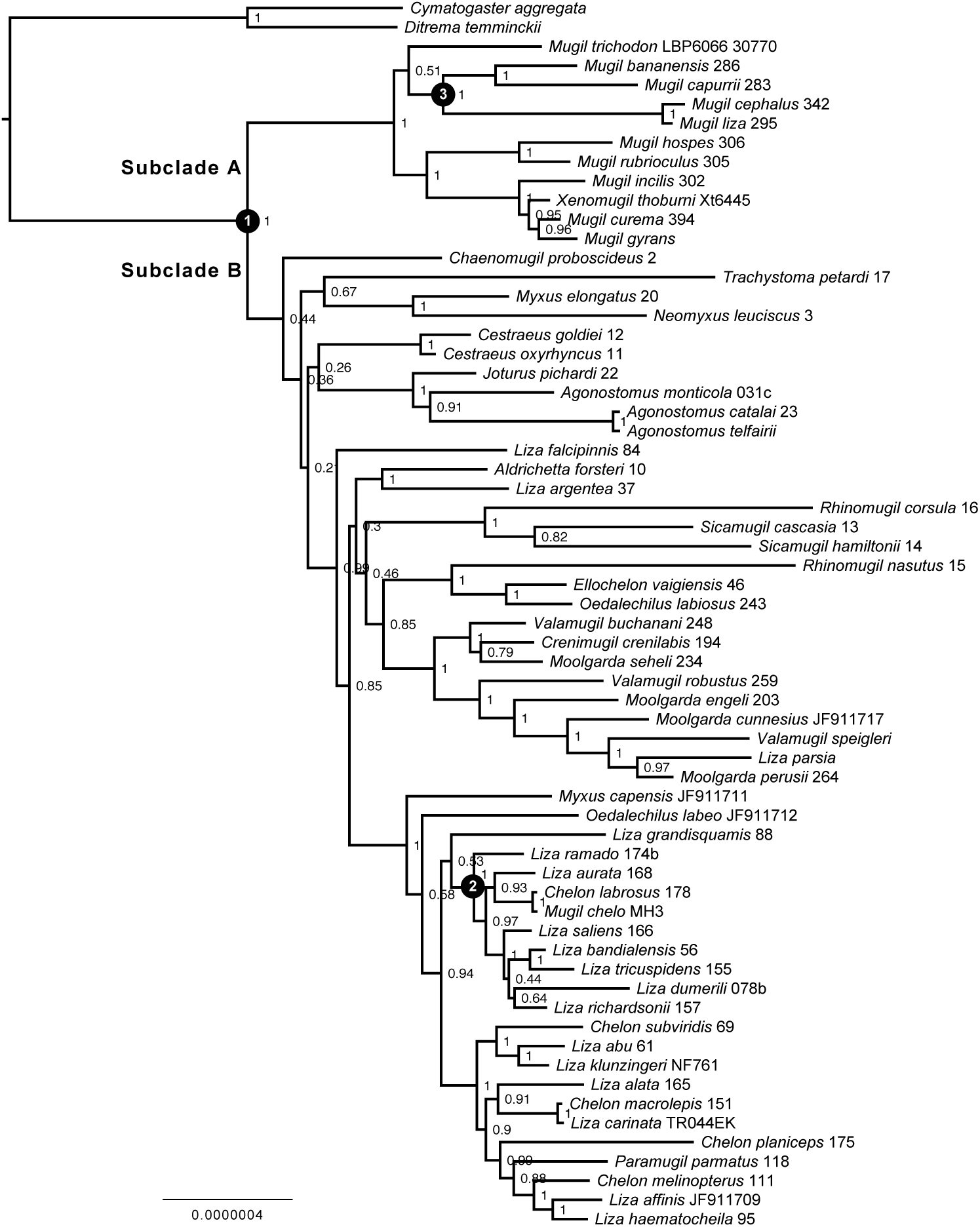
Bayesian estimate of mullet phylogeny. Estimates are based on the 64-species dataset under the preferred mixed-substitution model (PS2; Table S2). Branch lengths are rendered proportional to the expected number of substitutions per site (inset scale bar). Numbers adjacent to internal nodes indicate the corresponding marginal probabilities, and the three circled internal nodes indicate the location of the corresponding fossil calibrations (see Table 2).

### Divergence-time estimation

We evaluated six candidate relaxed-clock models to estimate divergence times for the 64-species dataset. These models comprise all possible combinations of the two mixed-substitution models (PS1 and PS2; Table S2), the three branch-rate models—the uncorrelated lognormal (UCLN) [41], uncorrelated exponential (UCEX) [41], and random-local molecular clock (RLC) [42] models—and the single node-age model—the sampled birth-death (SBD) model [43]. To render estimates in absolute time, we employed the same set of fossil calibrations as those used in the analyses of the 100-species dataset (Table 2). The results of our preliminary phylogenetic analyses indicate strong support (*P* ∼ 1.0) for all three prospective calibration points (Figure S6), however, we only constrained the monophyly on the ingroup node (calibration node 1).

We estimated the joint posterior probability distribution of the phylogeny, divergence times and other parameters under each of the six candidate relaxed-clock models using the MCMC algorithms implemented in `BEAST` v.1.8.2 [34]. For each relaxed-clock model, we ran four replicate MCMC simulations for 400 million cycles, thinned chains by sampling every 40, 000^th^ state, and assessed the reliability of the approximations. We then combined the stationary samples from the four independent simulations under each candidate relaxed-clock model.

We used the posterior samples for each of the six relaxed-clock models to assess their fit to the 64-species dataset using Bayes factors. Specifically, we estimated the marginal likelihood for each relaxed-clock model using the AICm method-of-moments estimator [63, 64, 68] implemented in `Tracer` v.1.6 [34]. Finally, we used these marginal-likelihood estimates (Table S4) to select among the corresponding relaxed-clock models using Bayes factors (Table S5).

**Table S4.** Marginal likelihoods of relaxed-clock models for the 64-species dataset. Model comparisons are based on analyses of the 64-species dataset. Marginal likelihoods for each of the candidate relaxed-clock models are based on the AICm method-of-moments estimator [63, 64]. Estimates of the standard error (S.E.) are based on 1000 bootstrap replicates performed in `Tracer` v.1.6.

| Relaxed-clock | Marginal likelihood |  |
| --- | --- | --- |
| Model | Mean | S.E. |
| RLC-PS1 | -68993.05 | ±0.85 |
| RLC-PS2 | -67503.97 | ±0.68 |
| UCEX-PS1 | -66440.25 | ±0.25 |
| UCEX-PS2 | -64889.63 | ±0.32 |
| UCLN-PS1 | -66437.12 | ±0.32 |
| UCLN-PS2 | -64884.17 | ±0.29 |

We summarized the resulting composite marginal posterior probability density for the preferred relaxed-clock model (UCLN+PS2+SBD) as a maximum clade credible (MCC) consensus tree with median node ages (Figure S7). We also explored the impact of the relaxed-clock models on estimated divergence times by comparing the inferred ages of four key nodes: (1) the mugilid stem age; (2) the mugilid crown age; (3) the crown age of Subclade A, and; (4) the crown age of Subclade B (Table S6). Clearly, the relaxed-clock model has a strong impact on divergence-time estimates, and the different model components differ in their influence. For a given branch-rate model, age estimates for alternate partition schemes differed on average by 7.5%, whereas for a given partition scheme, ages for alternative branch-rate models differed on average by 27.3%.

**Table S5.** Bayes factor comparisons of relaxed-clock models for the 64-species dataset. Model comparisons are based on analyses of the 64-species dataset. For each model comparison, *M*_0_ : *M*_1_, we calculated the Bayes factor as 2*ln*(*M*_0_ *− M*_1_). The table compares marginal likelihoods for the pair of models in row *i* and column *j*: positive values indicate support for the corresponding model in row *i*. The UCLN+PS2+SBD relaxed-clock model is strongly preferred over rival models (2.3 *> ln*BF > 4.6) [65].

| Relaxed-clock |  | Bayes Factor |  |  |  |  |
| --- | --- | --- | --- | --- | --- | --- |
| Model | RLC-PS1 | RLC-PS2 | UCEX-PS1 | UCEX-PS2 | UCLN-PS1 | UCLN-PS2 |
| RLC-PS1 | — | -1489.08 | -2552.80 | -4103.42 | -2555.93 | -4108.88 |
| RLC-PS2 | 1489.08 | — | -1063.73 | -2614.34 | -1066.85 | -2619.80 |
| UCEX-PS1 | 2552.80 | 1063.73 | — | -1550.61 | -3.12 | -1556.07 |
| UCEX-PS2 | 4103.42 | 2614.34 | 1550.61 | — | 1547.49 | -5.46 |
| UCLN-PS1 | 2555.93 | 1066.85 | 3.12 | -1547.49 | — | -1552.95 |
| UCLN-PS2 | 4108.88 | 2619.80 | 1556.07 | 5.46 | 1552.95 | — |

**Table S6.** The impact of relaxed-clock models on divergence-time estimates. Comparisons are based on analyses of the 64-species dataset. We report the estimated median and 95% HPD of ages for four key nodes under the six relaxed-clock models that we explored. ^†^The divergence-time estimates under the preferred relaxed-clock model.

| Node | Relaxed-Clock Model |  |  |  |  |  |
| --- | --- | --- | --- | --- | --- | --- |
|  | RLC-PS1 | RLC-PS2 | UCEX-PS1 | UCEX-PS2 | UCLN-PS1 | UCLN-PS2<sup>†</sup> |
| Mullet stem | 38.1 | 58.2 | 54.0 | 56.1 | 75.3 | 76.7 |
|  | [31.6,46.1] | [43.7,74.6] | [38.8,75.7] | [40.6,77.7] | [52.3,103.8] | [53.6,107.2] |
| Mullet crown | 36.0 | 39.8 | 48.3 | 50.6 | 63.5 | 65.0 |
|  | [30.9,42.7] | [33.1.,48.7] | [36.7,63.8] | [38.5,66.3] | [49.1,81.0] | [49.6,83.1] |
| Subclade A | 30.6 | 33.2 | 29.8 | 32.9 | 38.1 | 40.9 |
|  | [26.9,35.8] | [28.3,39.4] | [19.2,43.6] | [21.8,46.9] | [27.4,51.4] | [29.1,54.4] |
| Subclade B | 26.2 | 27.9 | 45.4 | 46.6 | 59.1 | 59.0 |
|  | [23.2,30.4] | [23.8,33.3] | [34.9,59.0] | [36.2,60.5] | [46.0,74.5] | [45.8,74.7] |

**Figure S7.**
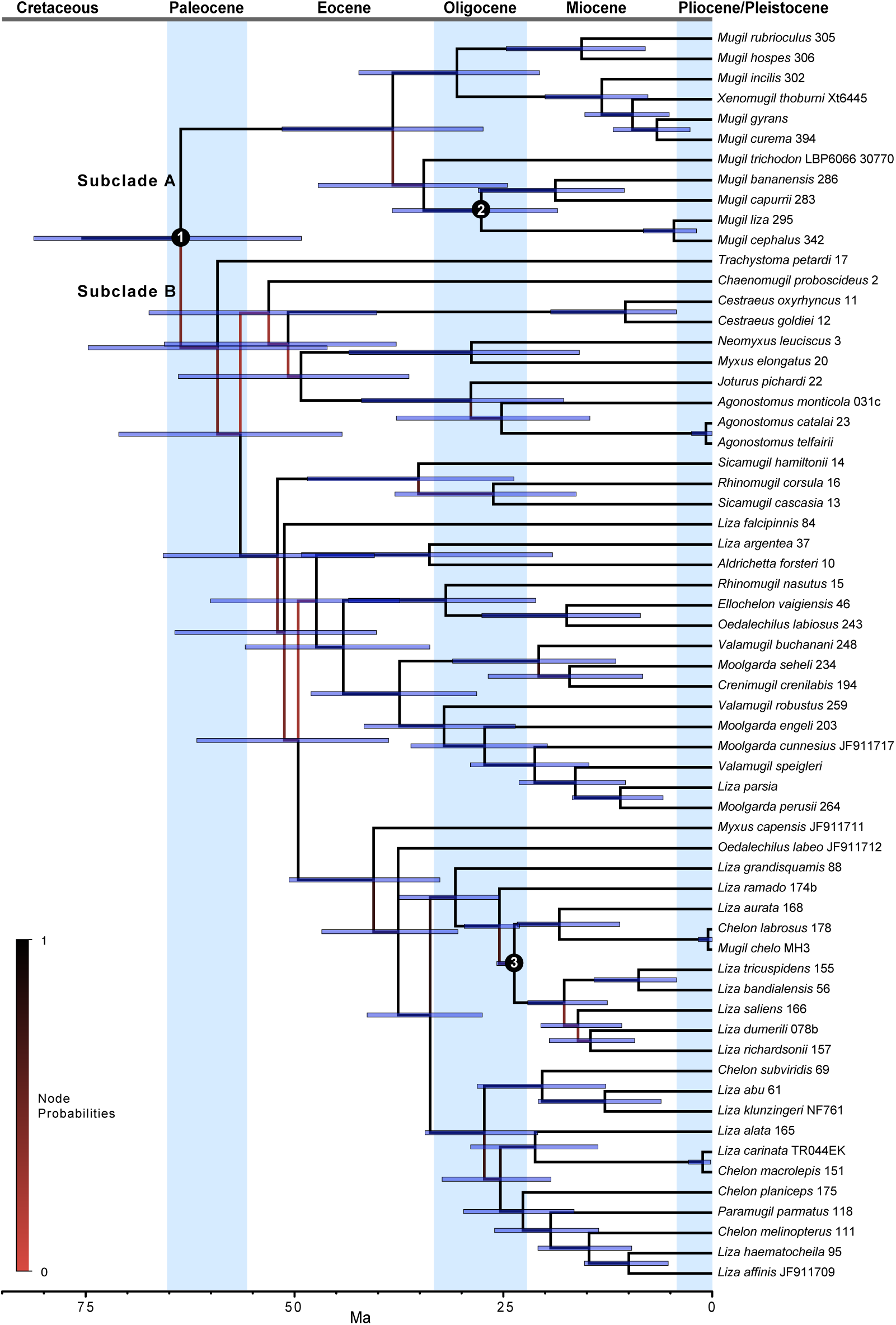
Bayesian estimate of mullet divergence times. Estimates are based on the 64-species dataset under the preferred relaxed-clock model (UCLN+PS2+SBD). The shading of internal branches indicates the corresponding node probabilities (see inset legend), the numbered internal nodes indicate the location of the corresponding fossil calibrations (Table 2), and the bars on nodes indicate the corresponding 95% HPD interval of divergence times.

### Ancestral-state estimation

We used the inferred phylogeny as a framework for exploring the evolution of feeding preference in mullets. We scored feeding preference as a discrete variable with three states (algae, detritus, or invertebrates), reflecting the main food item of each mullet species. We assessed the fit of these discrete traits to four candidate models, comprising all possible combinations of two continuous-time Markov (CTM) models—the first assumes a single instantaneous rate of change between the three discrete states (CTM-3), and the second assumes six instantaneous-rate parameters to describe changes between the three states (CTM-6)—and two branch-rate models—the first assumes that the rate of trait evolution is constant across branches (the continuous-rate morphological clock model, CRMC), and the second assumes that rates of trait evolution and substitution vary across branches under a shared model (the uncorrelated lognormal model, UCLN).

We conditioned inferences of diet evolution on the previously inferred MCC topology (Figure S7), but integrated out uncertainty in divergence times under the preferred relaxed-clock model (PS2+UCLN+SBD; Table S5) and the three fossil calibration densities (Table 2). We assumed uniform priors, Uniform[0, 1], for both the stationary and root frequencies of the three discrete states, and a mean-one gamma prior, Gamma[1, 1], on the instantaneous-rate parameters. We simultaneously estimated the number of changes in feeding preference in mullets—between diets of algae, detritus, or invertebrates—using the robust Markov-jump approach [66, 67] implemented in `BEAST` v.1.8.2 [34].

For each candidate discrete-trait model, we inferred the joint posterior probability by performing four replicate MCMC simulations of 400 million cycles in `BEAST` v.1.8.2 [34], thinning the chain by sampling every 4000^th^, and assessed the reliability of the approximations as previously. We combined the stationary samples from the four replicate simulations under each model, and used these composite posterior samples to assess the fit of the discrete-trait data to the the four candidate models. Specifically, we estimated the marginal likelihood for each discrete-trait model using the AICm method-of-moments estimator [63, 64, 68] implemented in `Tracer` v.1.6 [34]. We then used these marginal-likelihood estimates to select among the corresponding discrete-trait models using Bayes factors (Table S7). Finally, we plotted the marginal probabilities for diet on the internal nodes of the MCC consensus tree using `FigTree` v.1.4.2 (Figure S8), and summarized the instantaneous rates and number of changes between states (Table S8).

There is strong support for asymmetric rates of change among states: when the branch-rate model is held constant, the *ln*BF favor the CTM-6 rate model over the symmetric CTM-3 rate by a factor of 8.1 (CRMC) and 6.4 (UCLN). Similarly, there is strong support for clock-like rates of morphological change: when the CTM model is held constant, the *ln*BF favors the constant-rate morphological-clock model by a factor of 5.3 (CTM-3 rate) and 7.1 (CTM-6 rate) (see Table S7). This result suggests that—for the 64-species mullet dataset—rates of morphological evolution are not correlated with rates of substitution. It is also interesting to note that species sampling does not appear to strongly impact inferences of trait evolution: inferred ancestral states and the number of changes between states are very similar for the 100- and 64-species datasets (compare Figures 2–S8; Tables 9–S8).

**Table S7.** Marginal likelihoods and Bayes factor comparisons for discrete-trait models. Marginal-likelihood estimates and model comparisons are based on analyses of the 64-species dataset. Candidate discrete-trait models comprise all combinations of branch-rate models—the constant-rate morphological clock (CRMC) and the uncorrelated-exponential relaxed-clock (UCEX) models—and two site models, where rates of change between the three discrete-traits are assumed to be symmetric (CTM3) or are allowed to be assymmetric (CTM6). Marginal likelihoods for each of the candidate discrete-trait models are based on the AICm method-of-moments estimator [63, 64]. Estimates of the standard error (S.E.) are based on 1000 bootstrap replicates performed in `Tracer` v.1.6. For each model comparison, *M*_0_ : *M*_1_, we calculated the Bayes factor as 2*ln*(*M*_0_ *− M*_1_). The table compares marginal likelihoods for the pair of models in row *i* and column *j*: positive values indicate support for the corresponding model in row *i*. The CRMC-CTM 6-rate discrete-trait model is strongly preferred over competing models (2.3 *> ln*BF > 4.6) [65].

| Discrete-trait<br>Model | Marginal likelihood |  | Bayes Factor |  |  |  |
| --- | --- | --- | --- | --- | --- | --- |
|  | Mean | S.E. | CRMC-CTM3 | CRMC-CTM6 | UCLN-CTM3 | UCLN-CTM6 |
| CRMC-CTM3 | -66546.51 | ±0.38 | — | -8.14 | 5.29 | 0.00 |
| CRMC-CTM6 | -66538.37 | ±0.20 | 8.14 | — | 13.43 | 7.08 |
| UCLN-CTM3 | -66551.80 | ±0.29 | -5.29 | -13.43 | — | -6.35 |
| UCLN-CTM6 | -66545.45 | ±0.15 | 1.06 | -7.08 | 6.35 | — |

**Table S8.** Inferred rates and counts of diet change in mullets. We inferred the evolution of diet in mullets for the 64-species dataset under the preferred dicrete-trait model, which specifies that rates of diet evolution are constant across branches of the tree, and rates of change between the three discrete states are independent. Here we report the mean [and 95% HPD] of estimated instantaneous rates of change, *q*_*ij*_, between the three states—algae (A), detritus (D), and invertebrates (I)—and the expected number of changes between states estimated using the Markov-jump approach [66, 67].

| Instantaneous-rate parameter | Mean<br>rate | 95% HPD<br>rate | Mean<br>count | 95% HPD<br>count |
| --- | --- | --- | --- | --- |
| $q_{AD}$ | 0.76 | [6.98E-5,1.90] | 2.23 | [0.87, 4.54] |
| $q_{DA}$ | 0.53 | [3.56E-5,1.47] | 0.51 | [2.97E-6, 1.52] |
| $q_{AI}$ | 0.31 | [1.18E-5,0.99] | 1.24 | 5.89E-6, 3.22] |
| $q_{IA}$ | 0.38 | [3.28E-5,1.17] | 8.38 | [5.25, 12.16] |
| $q_{ID}$ | 1.50 | [2.38E-1,3.15] | 13.71 | [10.93, 16.31] |
| $q_{DI}$ | 2.37 | [5.58E-1,4.66] | 0.30 | [1.75E-6, 1.17] |

**Figure S8.**
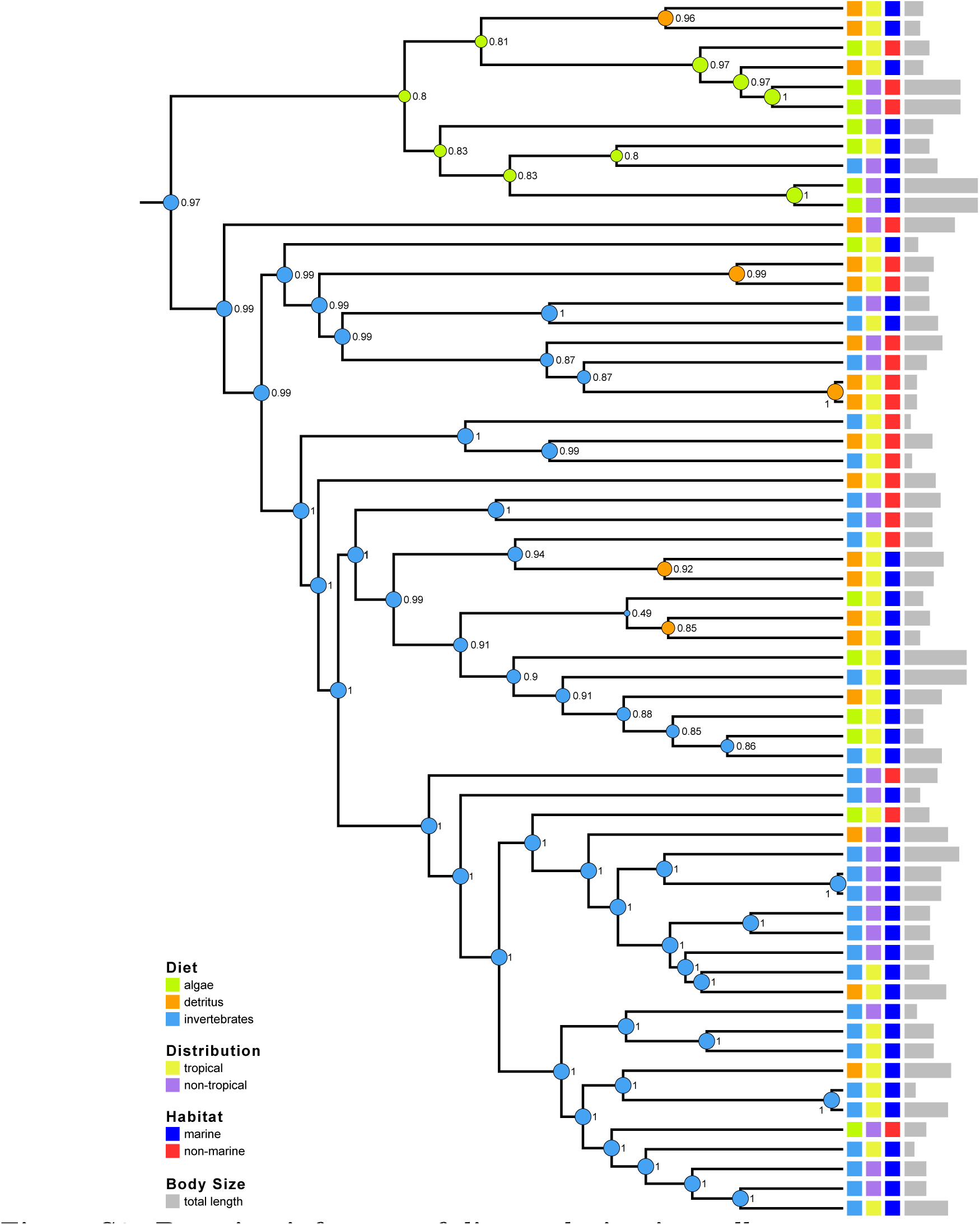
Bayesian inference of diet evolution in mullets. Estimates are based on the 64-species dataset under the preferred discrete-trait model (CRMC-CTM6). Circles at interior nodes are colored according to the MAP estimate of the ancestral diet—algae, detritus, or invertebrates—where the diameter and adjacent numbers indicate the marginal posterior probability of the MAP state. Other variables—biogeographic distribution, habitat, and body size—are indicated at the tips of the tree for each species (see inset legend).

### Correlated-trait evolution

We explored correlations between traits in the 64-species tree using the latent-liability model [69]. The four traits included two continuous and two discrete variables: trophic index (TI, continuous), total length (TL, continuous), habitat (Hab, discrete), and geographic distribution (Dist, discrete). The latent-liability model assumes continuous traits can realize any positive or negative value; therefore, it was necessary to transform our continuous traits to satisfy this assumption. Specifically, we normalized trophic-index values (which range from 2.0 to 3.4) so that they ranged from 0 and 1, and then logit-transformed them; the logit-transformed trophic-index values range from − ∞ to ∞. We also ln-transformed total-length values (which range from 0 to ∞), resulting in values between − ∞.and ∞. We treated both discrete traits as binary; habitat was scored as ‘marine’ or ‘non-marine’, and distribution was scored as ‘tropical’ or ‘non-tropical’.

We analyzed our trait data on a fixed tree (the MCC phylogeny estimated above) using the latent-liability module [69] implemented in `BEAST` v. 1.8.3 [34]. To assess the sensitivity of our analysis to prior specification, we explored three different values for the rate-matrix parameter **R** (low, medium, and high), as well as three different prior values for the precision-matrix parameter (low, medium, and high), and used a fixed value of ***d*** = 6 for all analyses. For each combination of prior values (9 in total), we ran four independent MCMC simulations for 200 million generations, thinning each chain by sampling every 20,000^th^ state. We assessed the performance of each MCMC simulation in the usual manner using `Tracer` [34] and `coda` [35]. We then discarded the first 25% of samples from each simulation, and combined the stationary samples from each of the four runs, providing 30,000 samples from which to estimate the marginal posterior densities of covariances among traits for each of the nine prior combinations.

Finally, we transformed the marginal densities of evolutionary covariances into marginal densities of correlation coefficients, which range from −1 to 1 to provide a more natural interpretation of evolutionary correlation that can be compared among traits, regardless of the overall rate of evolution. For each marginal density, we identified the correlation coefficient as significantly different from zero if a correlation coefficient of zero (*i.e.*, no correlation) was as or more extreme than 95% of the marginal density. Across all prior combinations, total length and distribution were the only significant correlations (Figure S9). The estimated correlation coefficients were qualitatively identical among each of the prior combinations, indicating that our results are robust to prior specification.

**Figure S9.**
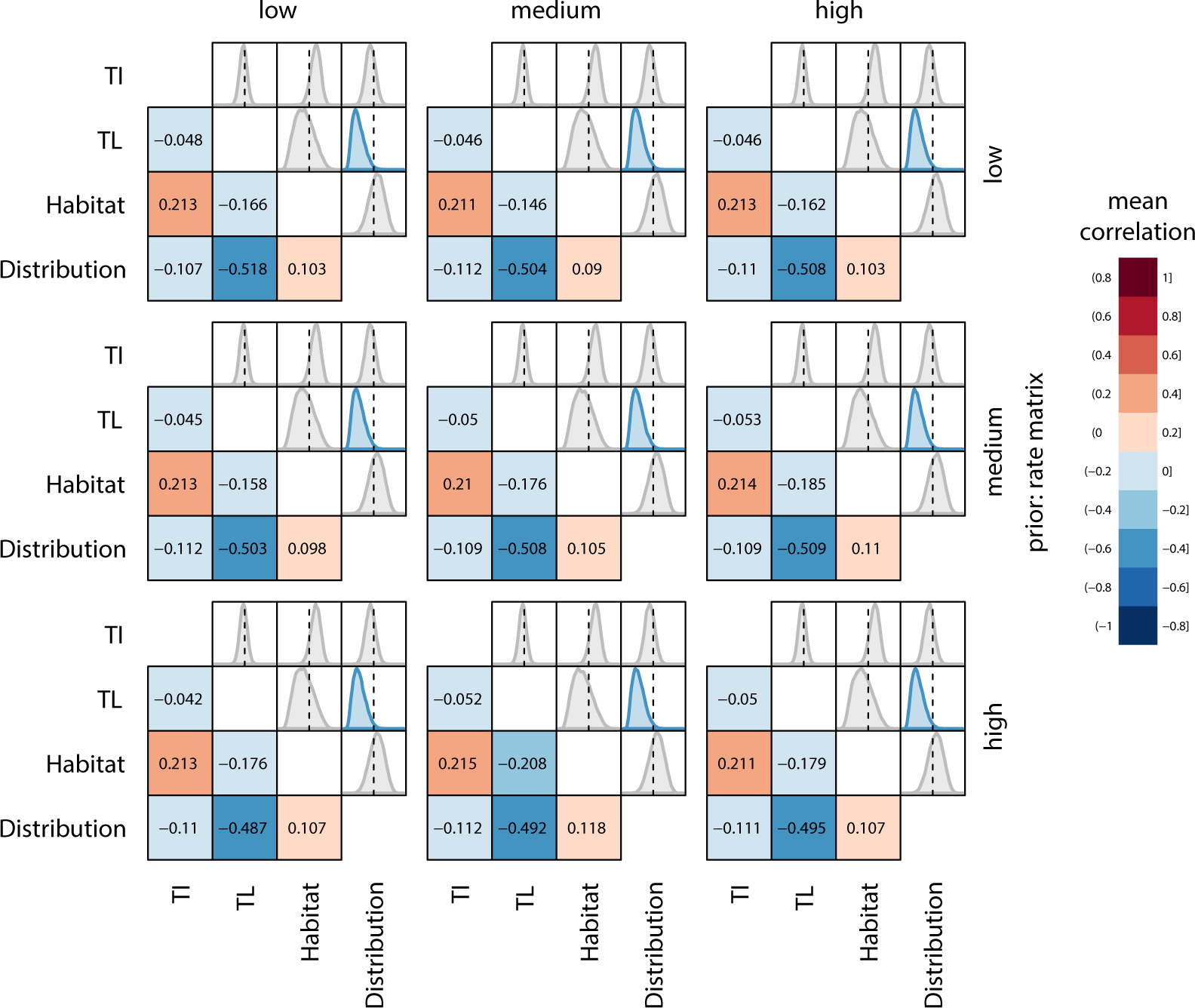
Correlates of diet evolution in mullets. Estimates are based on analyses of the 64-species dataset under the latent-liability model [69]. Traits (abbreviations) [and states] include: trophic index (TI); total length (TL) [centimeters]; habitat type (Habitat) [marine, non-marine]; and distribution (Distribution) [tropical, non-tropical]. We repeated these analyses using nine distinct combinations of priors on the precision-matrix and rate-matrix parameters. Each row of panels corresponds to low, medium, or high values for the rate-matrix parameter; each column corresponds to low, medium and high values for the precision-matrix parameter. Within each panel, the lower diagonal depcts the mean correlation coefficients for each pair of traits (see inset legend), and the upper diagonal depicts the corresponding marginal densities of the correlation coefficients (values range from −1 to +1). Densities are colored according to their mean value only if they differ significantly from zero (*i.e.*, the posterior probability that the value is equal to or more extreme than 0 is < 0.05).

**Table S9.** Sequence data used in study of mullet phylogeny.

| Species | Isolate | 16S | COI | cytb |
| --- | --- | --- | --- | --- |
| *Cymatogaster aggregata* | NC 009059 | AP009128 | AP009128 | AP009128 |
| *Ditrema temminckii* | NC 009060 | AP009129 | AP009129 | AP009129 |
| *Agonostomus catalai* | 23 | JQ060643 | JQ060394 | JQ060138 |
| *Agonostomus monticola* | 26 | JQ060645 | JQ060395 | JQ060139 |
| *Agonostomus monticola* | 35 | JQ060644 | JQ060403 | JQ060147 |
| *Agonostomus monticola* | AP002930<sup>†</sup> | JF911702 | JF911702 | JF911702 |
| *Aldrichetta forsteri* | 10 | JQ060654 | JQ060405 | JQ072905 |
| *Aldrichetta forsteri* | JF911703<sup>†</sup> | JF911703 | JF911703 | JF911703 |
| *Cestraeus goldiei* | 12 | JQ060655 | JQ060406 | JQ060149 |
| *Cestraeus oxyrhyncus* | 11 | JQ060656 | JQ060407 | JQ060150 |
| *Chaenomugil proboscideus* | 2 | JQ060657 | JQ060408 | JQ060151 |
| *Chelon labrosus* | 179b | JQ060660 | JQ060411 | JQ060154 |
| *Chelon macrolepis* | 124 | JQ060662 | JQ060414 | JQ060157 |
| *Chelon macrolepis* | 146 | JQ060670 | JQ060424 | JQ060167 |
| *Chelon melinopterus* | 111 | JQ060676 | JQ060428 | JQ060171 |
| *Chelon planiceps* | 122 | JQ060679 | JQ060429 | JQ060172 |
| *Chelon planiceps* | 175 | JQ060677 | JQ060430 | JQ060173 |
| *Chelon subviridis* | 64 | JQ060680 | JQ060432 | JQ060175 |
| *Chelon subviridis* | 69 | JQ060682 | JQ060433 | JQ060176 |
| *Chelon subviridis* | 70 | JQ060681 | JQ060434 | JQ060177 |
| *Crenimugil crenilabis* | 187 | JQ060683 | JQ060436 | JQ060179 |
| *Ellochelon vaigiensis* | 46 | JQ060691 | JQ060443 | JQ060186 |
| *Ellochelon vaigiensis* | 51 | JQ060692 | JQ060444 | JQ060187 |
| *Joturus pichardi* | 22 | JQ060694 | JQ060446 | JQ060189 |
| *Liza abu* | 61 | JQ060695 | JQ060447 | JQ060190 |
| *Liza affinis* | 057x | JQ060696 | JQ060448 | JQ060191 |
| *Liza alata* | 162 | JQ060697 | JQ060449 | JQ060192 |
| *Liza alata* | 165b | JQ060700 | JQ060452 | JQ060195 |
| *Liza argentea* | 39 | JQ060702 | JQ060454 | JQ060197 |
| *Liza aurata* | 171c | JQ060706 | JQ060459 | JQ060202 |
| *Liza bandialensis* | 54 | JQ060708 | JQ060461 | JQ060204 |
| *Liza carinata* | TR044EK | – | JQ623947 | – |
| *Liza dumerili* | 76 | JQ060711 | JQ060464 | JQ060207 |
| *Liza dumerili* | 79 | JQ060714 | JQ060467 | JQ060210 |
| *Liza falcipinnis* | 83 | JQ060716 | JQ060469 | JQ060212 |
| *Liza grandisquamis* | 91 | JQ060723 | JQ060476 | JQ060219 |
| *Liza haematocheila* | 102 | JQ060725 | JQ060478 | JQ060221 |
| *Liza klunzingeri* | NF762 | – | JX983356 | – |
| *Liza parsia* | 062b | JQ060739 | JQ060493 | JQ060237 |
| *Liza ramado* | 173 | GQ258707 | JQ060479 | JQ060222 |
| *Liza richardsonii* | 157 | JQ060727 | JQ060481 | JQ060224 |
| *Liza saliens* | 166 | GQ258709 | JQ060483 | JQ060226 |
| *Liza tricuspidens* | 155 | JQ060740 | JQ060495 | JQ060238 |
| *Moolgarda cunnesius* | 275 | JQ060743 | JQ060496 | JQ060239 |
| *Moolgarda cunnesius* | 276 | JQ060742 | JQ060497 | JQ060240 |
| *Moolgarda cunnesius* | 276b | JQ060744 | JQ060499 | JQ060242 |
| *Moolgarda cunnesius* | 278 | JQ060741 | JQ060498 | JQ060241 |
| *Moolgarda engeli* | 203 | JQ060748 | JQ060503 | JQ060246 |
| *Moolgarda perusii* | 264 | JQ060749 | JQ060504 | JQ060247 |
| *Moolgarda perusii* | 274 | JQ060750 | JQ060505 | JQ060248 |
| *Moolgarda seheli* | 210 | JQ060755 | JQ060510 | JQ060253 |
| *Moolgarda seheli* | 217 | JQ060756 | JQ060511 | JQ060254 |
| *Moolgarda seheli* | 220 | JQ060758 | JQ060513 | JQ060256 |
| *Moolgarda seheli* | 225 | JQ060759 | JQ060514 | JQ060257 |
| *Moolgarda seheli* | 226 | JQ060760 | JQ060515 | JQ060258 |
| *Moolgarda seheli* | 235 | JQ060763 | JQ060517 | JQ060261 |
| *Mugil bananensis* | 286 | JQ060769 | JQ060523 | JQ060267 |
| *Mugil capurrii* | 283 | HM143895 | JQ060526 | JQ060270 |
| *Mugil cephalus* | AP002930<sup>†</sup> | AP002930 | AP002930 | AP002930 |
| *Mugil cephalus* | 325c | JQ060819 | JQ060536 | JQ060280 |
| *Mugil cephalus* | 326 | JQ060796 | JQ060537 | JQ060281 |
| *Mugil cephalus* | 329b | JQ060778 | JQ060541 | JQ060285 |
| *Mugil cephalus* | 342 | JQ060810 | JQ060545 | JQ060289 |
| *Mugil cephalus* | 344 | JQ060786 | HQ149711 | JQ060293 |
| *Mugil cephalus* | 349 | JQ060774 | HQ149714 | JQ060295 |
| *Mugil cephalus* | 350 | JQ060820 | JQ060549 | JQ060296 |
| *Mugil cephalus* | 358 | JQ060803 | JQ060553 | JQ060300 |
| *Mugil cephalus* | 361 | JQ060804 | JQ060554 | JQ060301 |
| *Mugil cephalus* | 368 | JQ060785 | JQ060559 | JQ060306 |
| *Mugil cephalus* | 375 | JQ060790 | Q060563 | JQ060311 |
| *Mugil cephalus* | 377 | JQ060814 | JQ060565 | JQ060313 |
| *Mugil curema* | 390 | JQ060843 | JQ060575 | JQ060324 |
| *Mugil curema* | 403 | JQ060841 | JQ060585 | JQ060334 |
| *Mugil curema* | 405 | JQ060823 | JQ060587 | JQ060336 |
| *Mugil curema* | 406 | JQ060830 | JQ060588 | JQ060337 |
| *Mugil curema* | 423 | JQ060835 | JQ060597 | JQ060346 |
| *Mugil hospes* | 306 | JQ060857 | JQ060607 | JQ060356 |
| *Mugil hospes* | LBP6061 42661 | JX000561 | JX185218 | JX185297 |
| *Mugil incilis* | 299 | JQ060859 | JQ060609 | JQ060358 |
| *Mugil liza* | 298 | JQ060861 | HQ149713 | JQ060360 |
| *Mugil rubrioculus* | 305 | JQ060864 | JQ060612 | JQ060363 |
| *Mugil rubrioculus* | 305b | JQ060865 | JQ060613 | JQ060364 |
| *Mugil rubrioculus* | LBP9060 42655 | JX000555 | JX185212 | JX185291 |
| *Mugil trichodon* | 291 | JQ060866 | JQ060614 | JQ060365 |
| *Myxus capensis* | 19 | JQ060867 | JQ060615 | JQ060366 |
| *Myxus elongatus* | 20 | JQ060868 | JQ060616 | JQ060367 |
| *Neomyxus leuciscus* | 3b | JQ060870 | JQ060618 | JQ060369 |
| *Oedalechilus labeo* | 181 | JQ060871 | JQ060619 | JQ060370 |
| *Oedalechilus labiosus* | 243 | JQ060872 | JQ060620 | JQ060371 |
| *Paramugil parmatius* | 118 | JQ060873 | JQ060621 | JQ060372 |
| *Rhinomugil corsula* | 16 | JQ060874 | JQ060622 | JQ060373 |
| *Rhinomugil nasutus* | 15 | JQ060875 | JQ060623 | JQ060374 |
| *Sicamugil cascasia* | JF911715<sup>†</sup> | JF911715 | JF911715 | JF911715 |
| *Sicamugil hamiltonii* | 14 | JQ060877 | JQ060625 | JQ060376 |
| *Trachystoma petardi* | 17 | JQ060878 | JQ060626 | JQ060377 |
| *Valamugil buechanani* | 245 | JQ060879 | JQ060627 | JQ06037B |
| *Valamugil buechanani* | 248 | JQ060894 | JQ060641 | JQ060392 |
| *Valamugil robustus* | 259 | JQ060881 | JQ060629 | JQ060380 |
| *Valamugil speigleri* | Os Vs 2 | KF375073 | JQ045778 | KF375151 |
| *Xenomugil thoburni* | XT6445 | JX559530 | JX559535 | JX559526 |

<sup>†</sup>individual gene regions excised from complete mitochondrial genome.

**Table S10.** Trait data used in study of feeding evolution. A (alage), D (detritus), I (invertebrates); M (marine), F (freshwater), B (brackish).

| Species | Diet | TROPH | Length | Habitat | Distribution |
| --- | --- | --- | --- | --- | --- |
| *Cymatogaster aggregata* | I | 3.0 | 20 | M | subtropical |
| *Ditrema temminckii* | I | 3.4 | 30 | M | temperate |
| *Agonostomus catalai* | D | 2.4 | 20 | F | tropical |
| *Agonostomus monticola* | I | 3.4 | 36 | F | subtropical |
| *Agonostomus monticola* | I | 3.4 | 36 | F | subtropical |
| *Agonostomus monticola* | I | 3.4 | 36 | F | subtropical |
| *Aldrichetta forsteri* | I | 2.5 | 58 | B | temperate |
| *Aldrichetta forsteri* | I | 2.5 | 58 | B | temperate |
| *Cestraeus goldiei* | D | 2.4 | 47 | F | tropical |
| *Cestraeus oxyrhyncus* | D | 2.4 | 39 | F | tropical |
| *Chaenomugil proboscideus* | A | 2.0 | 22 | M | tropical |
| *Chelon labrosus* | I | 2.6 | 88 | M | subtropical |
| *Chelon macrolepis* | I | 2.6 | 70 | M | tropical |
| *Chelon macrolepis* | I | 2.6 | 70 | M | tropical |
| *Chelon melinopterus* | D | 2.3 | 35 | M | subtropical |
| *Chelon planiceps* | A | 2.0 | 70 | M | tropical |
| *Chelon planiceps* | A | 2.0 | 70 | M | tropical |
| *Chelon subviridis* | I | 2.7 | 47 | M | tropical |
| *Chelon subviridis* | I | 2.7 | 47 | M | tropical |
| *Chelon subviridis* | I | 2.7 | 47 | M | tropical |
| *Crenimugil crenilabis* | D | 2.3 | 60 | M | tropical |
| *Ellochelon vaigiensis* | D | 2.3 | 63 | M | tropical |
| *Ellochelon vaigiensis* | D | 2.3 | 63 | M | tropical |
| *Joturus pichardi* | D | 2.4 | 61 | F | subtropical |
| *Liza abu* | I | 2.6 | 20 | F | subtropical |
| *Liza affinis* | A | 2.9 | 35 | B | subtropical |
| *Liza alata* | D | 2.3 | 75 | M | tropical |
| *Liza alata* | D | 2.3 | 75 | M | tropical |
| *Liza argentea* | I | 2.9 | 45 | B | temperate |
| *Liza aurata* | I | 2.5 | 59 | M | temperate |
| *Liza bandialensis* | I | 2.5 | 67 | M | tropical |
| *Liza carinata* | I | 2.6 | 18 | M | tropical |
| *Liza dumerili* | I | 2.7 | 40 | M | tropical |
| *Liza dumerili* | I | 2.7 | 40 | M | tropical |
| *Liza falcipinnis* | D | 2.3 | 50 | B | tropical |
| *Liza grandisquamis* | A | 2.0 | 40 | B | tropical |
| *Liza haematocheila* | I | 2.5 | 80 | B | tropical |
| *Liza klunzingeri* | I | 2.6 | 20 | M | subtropical |
| *Liza parsia* | A | 2.0 | 16 | M | tropical |
| *Liza ramado* | D | 2.2 | 70 | M | temperate |
| *Liza richardsonii* | D | 2.4 | 41 | M | subtropical |
| *Liza saliens* | I | 3.0 | 47 | M | subtropical |
| *Liza tricuspidens* | I | 2.5 | 75 | B | tropical |
| *Moolgarda cunnesius* | D | 2.4 | 41 | M | tropical |
| *Moolgarda cunnesius* | D | 2.4 | 41 | M | tropical |
| *Moolgarda cunnesius* | D | 2.4 | 41 | M | tropical |
| *Moolgarda cunnesius* | D | 2.4 | 41 | M | tropical |
| *Moolgarda engeli* | I | 2.5 | 30 | M | tropical |
| *Moolgarda perusii* | I | 2.5 | 25 | M | tropical |
| *Moolgarda perusii* | I | 2.5 | 25 | M | tropical |
| *Moolgarda seheli* | D | 2.3 | 60 | M | tropical |
| *Moolgarda seheli* | D | 2.3 | 60 | M | tropical |
| *Moolgarda seheli* | D | 2.3 | 60 | M | tropical |
| *Moolgarda seheli* | D | 2.3 | 60 | M | tropical |
| *Moolgarda seheli* | D | 2.3 | 60 | M | tropical |
| *Moolgarda seheli* | D | 2.3 | 60 | M | tropical |
| *Mugil bananensis* | I | 2.8 | 40 | M | tropical |
| *Mugil capurrii* | A | 2.0 | 53 | M | subtropical |
| *Mugil cephalus* | A | 2.1 | 118 | M | subtropical |
| *Mugil cephalus* | A | 2.1 | 118 | M | subtropical |
| *Mugil cephalus* | A | 2.1 | 118 | M | subtropical |
| *Mugil cephalus* | A | 2.1 | 118 | M | subtropical |
| *Mugil cephalus* | A | 2.1 | 118 | M | subtropical |
| *Mugil cephalus* | A | 2.1 | 118 | B | subtropical |
| *Mugil cephalus* | A | 2.1 | 118 | M | subtropical |
| *Mugil cephalus* | A | 2.1 | 118 | M | subtropical |
| *Mugil cephalus* | A | 2.1 | 118 | M | subtropical |
| *Mugil cephalus* | A | 2.1 | 118 | M | subtropical |
| *Mugil cephalus* | A | 2.1 | 118 | M | subtropical |
| *Mugil cephalus* | A | 2.1 | 118 | M | subtropical |
| *Mugil cephalus* | A | 2.1 | 118 | M | subtropical |
| *Mugil cephalus* | A | 2.1 | 118 | M | subtropical |
| *Mugil curema* | A | 2.0 | 90 | B | subtropical |
| *Mugil curema* | A | 2.0 | 90 | B | subtropical |
| *Mugil curema* | A | 2.0 | 90 | B | subtropical |
| *Mugil curema* | A | 2.0 | 90 | B | subtropical |
| *Mugil curema* | A | 2.0 | 90 | B | subtropical |
| *Mugil hospes* | D | 2.3 | 25 | M | tropical |
| *Mugil hospes* | D | 2.3 | 25 | M | tropical |
| *Mugil incilis* | A | 2.0 | 40 | B | tropical |
| *Mugil liza* | A | 2.0 | 80 | B | tropical |
| *Mugil rubrioculus* | D | 2.3 | 30 | M | tropical |
| *Mugil rubrioculus* | D | 2.3 | 30 | M | tropical |
| *Mugil rubrioculus* | D | 2.3 | 30 | M | tropical |
| *Mugil trichodon* | A | 2.0 | 46 | M | subtropical |
| *Myxus capensis* | I | 2.8 | 53 | F | subtropical |
| *Myxus elongatus* | I | 3.1 | 40 | M | temperate |
| *Neomyxus leuciscus* | I | 2.9 | 54 | M | tropical |
| *Oedalechilus labeo* | I | 2.5 | 25 | M | subtropical |
| *Oedalechilus labiosus* | D | 2.4 | 47 | M | tropical |
| *Paramugil parmatius* | I | 2.5 | 30 | M | tropical |
| *Rhinomugil corsula* | D | 2.4 | 45 | F | tropical |
| *Rhinomugil nasutus* | I | 2.9 | 45 | F | tropical |
| *Sicamugil cascasia* | I | 2.7 | 10 | F | tropical |
| *Sicamugil hamiltonii* | I | 2.7 | 12 | F | tropical |
| *Trachystoma petardi* | D | 2.3 | 81 | F | subtropical |
| *Valamugil buechanani* | A | 2.2 | 100 | M | tropical |
| *Valamugil buechanani* | A | 2.2 | 100 | M | tropical |
| *Valamugil robustus* | A | 2.0 | 30 | B | tropical |
| *Valamugil speigleri* | A | 2.2 | 35 | M | tropical |
| *Xenomugil thoburni* | D | 2.3 | 30 | M | tropical |

## References

1. Harrison IJ, Senou H (1999) The living marine resources of the western central Pacific, Vol. 4 Bony fishes part 2 (Mugilidae to Carangidae), Rome: Food and Agriculture Organization. pp. 2069–2083.

2. Thomson JM (1997) The Mugilidae of the world. Memoirs of the Queensland Museum 41: 457–562.

3. Harrison IJ (2002) The living marine resources of the western central Atlantic, Vol. 2: Bony fishes part 1 (Acipenseridae to Grammatidae), Rome: Food and Agriculture Organization, chapter Order Mugiliformes, Mugilidae. pp. 1071–1085.

4. Nelson JS (2006) Fishes of the world. New Jersey, USA: John Wiley Sons.

5. Lobato FL, Barneche DR, Siqueira AC, Liedke AMR, Lindner A, et al. (2014) Diet and diversification in the evolution of coral reef fishes. PLoS One 9: e102094.

6. Gerking SH (1994) Feeding Ecology of Fish. San Diego: Academic Press.

7. Harrison IJ, Howes GJ (1991) The pharyngobranchial organ of mugilid fishes: its structure, variability, ontogeny, possible function and taxonomic utility. Bulletin of the British Museum of Natural History (Zoology) 57: 111–132.

8. Bowen SH (1983) Detritivory in Neotropical fish communities. Environmental Biology of Fishes 9: 137–144.

9. Choat JH, Clements KD (1998) Vertebrate herbivores in marine and terrestrial environments: a nutritional ecology perspective. Annual Review of Ecology and Systematics 29: 375–403.

10. German DP, Gawlick AK, Horn M (2014) Evolution of ontogenetic dietary shifts and associated gut features in prickleback fishes (Teleostei: Stichaeidae). Comparative Biochemistry and Physiology, Part B 168: 12–18.

11. Berg LS (1940) Classification of fishes, both recent and fossil. J.W. Edwards.

12. Gosline WA (1971) Functional morphology and classification of teleostean fishes. Honolulu: University Press of Hawaii.

13. Johnson GD, Patterson C (1993) Percomorph phylogeny: a survey of acanthomorphs and a new proposal. Bulletin of Marine Science 52: 554–626.

14. Mabuchi K, Miya M, Azuma Y, Nishida M (2007) Independent evolution of the specialized pharyngeal jaw apparatus in cichlid and labrid fishes. BMC Evolutionary Biology 7: 10.

15. Santini F, Harmon LJ, Carnevale G, Alfaro ME (2009) Did genome duplication drive the origin of teleosts? a comparative study of diversification in ray-finned fishes. BMC Evol Biol 9: 194.

16. Wainwright PC, Smith WL, Price SA, Tang KL, Sparks JS, et al. (2012) The evolution of pharyngognathy: a phylogenetic and functional appraisal of the pharyngeal jaw key innovation in labroid fishes and beyond. Syst Biol 61: 1001–1027.

17. Betancur-R R, Broughton RE, Wiley EO, Carpenter K, López JA, et al. (2013) The tree of life and a new classification of bony fishes. PLoS Currents Tree of Life DOI: 10.1371/currents.tol.53ba26640df0ccaee75bb165c8c26288.

18. Near TJ, Dornburg A, Eytan RI, Keck BP, Smith WL, et al. (2013) Phylogeny and tempo of diversification in the superradiation of spiny-rayed fishes. Proc Natl Acad Sci U S A 110: 12738–12743.

19. Rabosky DL, Santini F, Eastman J, Smith SA, Sidlauskas B, et al. (2013) Rates of speciation and morphological evolution are correlated across the largest vertebrate radiation. Nat Commun 4: 1958.

20. Froese R, Pauly D. FishBase: World wide web electronic publication.

21. Durand JD, Shen KN, Chen WJ, Jamandre B, Blel H, et al. (2012) Systematics of the grey mullets (Teleostei: Mugiliformes: Mugilidae): Molecular phylogenetic evidence challenges two centuries of morphology-based taxonomy. Molecular Phylogenetics and Evolution 64: 73–92.

22. Aurelle D, Barthelemy RM, Quignard JP, Trabelsi M, Faure E (2008) Molecular phylogeny of Mugilidae (Teleostei: Perciformes). The Open Marine Biology Journal 2: 29–37.

23. Heras S, Roldan MI, Castro MG (2009) Molecular phylogeny of Mugilidae fishes revised. Reviews in Fish Biology and Fisheries 19: 217–231.

24. Durand JD, Chen WJ, Shen KN, Fu C, Borsa P (2012) Genus-level taxonomic changes implied by the mitochondrial phylogeny of grey mullets (Teleostei: Mugilidae). Comptes Rendus Biologies 335: 687–697.

25. McMahan CD, Davis MP, Domnguez-Domnguez O, Garca-de Leon FJ, Doadrio I, et al. (2013) From the mountains to the sea: phylogeography and cryptic diversity within the mountain mullet, *Agonostomus monticola* (Teleostei: Mugilidae). Journal of Biogeography 40: 894–904.

26. Nematzadeh M, Gillkolaei S, Khalesi M, Laloei F (2013) Molecular phylogeny of mullets (Teleosti: Mugilidae) in Iran based on mitochondrial DNA. Biochemical Genetics 51: 334–340.

27. Edgar RC (2004) MUSCLE: multiple sequence alignment with high accuracy and high throughput. Nucleic Acids Research 32: 1792–1797.

28. Larsson A (2014) AliView: a fast and lightweight alignment viewer and editor for large datasets. Bioinformatics 30: 3276–3278.

29. Briggs JC, Bowen B (2012) A realignment of marine biogeographic provinces with particular reference to fish distributions. Journal of Biogeography 39: 12–30.

30. Zhang J, Kapli P, Pavlidis P, Stamatakis A (2013) A general species delimitation method with applications to phylogenetic placements. Bioinformatics 29: 2869–2876.

31. Lanfear R, Calcott B, Ho SYW, Guindon S (2012) PartitionFinder: Combined selection of partitioning schemes and substitution models for phylogenetic analyses. Molecular Biology and Evolution 29: 1695–1701.

32. Ronquist F, Teslenko M, van der Mark P, Ayres DL, Darling A,et al. (2012) MrBayes 3.2: Effcient Bayesian phylogenetic inference and model choice across a large model space. Systematic Biology 61: 539–542.

33. Schwarz G, et al. (1978) Estimating the dimension of a model. The annals of statistics 6: 461–464.

34. Drummond AJ, Suchard MA, Xie D, Rambaut A (2012) Bayesian phylogenetics with BEAUti and the BEAST 1.7. Molecular Biology And Evolution 29: 1969–1973.

35. Plummer M, Best N, Cowles K, Vines K (2006) CODA: Convergence diagnosis and output analysis for MCMC. R News 6: 7–11.

36. Brooks SP, Gelman A (1997) General methods for monitoring convergence of iterative simulations. Journal of Computational and Graphical Statistics 7: 434–455.

37. Geweke J (1992) Evaluating the accuracy of sampling-based approaches to the calculation of posterior moments (with discussion). In: Bernardo J, Berger J, Dawid A, Smith A, editors, Bayesian Statistics 4, Oxford: Oxford University Press. pp. 169–193.

38. Gelman A, Rubin D (1992) Inference from iterative simulation using multiple sequences (with discussion). Statistical Science 7: 457–511.

39. Heath TA, Moore BR (2014) Bayesian inference of species divergence times. In: Ming-Hui Chen LK, Lewis P, editors, Bayesian Phylogenetics: Methods, Algorithms, and Applications. Sinauer Associates, Sunderland, MA., pp. 487–533.

40. Akaike H (1974) A new look at the statistical model identification. Automatic Control, IEEE Transactions on 19: 716–723.

41. Drummond A, Ho S, Phillips M, Rambaut A (2006) Relaxed phylogenetics and dating with confidence. PLoS Biol 4: e88.

42. Drummond AJ, Suchard MA (2010) Bayesian random local clocks, or one rate to rule them all. BMC Biology 8: 114.

43. Gernhard T (2008) The conditioned reconstructed process. Journal of Theoretical Biology 253: 769–778.

44. Yule GU (1924) A mathematical theory of evolution, based on the conclusions of Dr. J. C. Wills, F. R. S. Philosophical Transactions of the Royal Society of London, Biology 213: 21–87.

45. Kendall DG (1948) On the generalized“birth-and-deat” process. The Annals of Mathematical Statistics 19: 1–15.

46. Switshenskaja AA (1973) Iskopemye kefalobraznye SSSR. Trudy Paleontologicheskogo Instituta Akademii Nauk SSSR 138: 1–64 [in Russian].

47. Trewavas E, Ingham SE (1972) A key to the species of Mugilidae (Pisces) in the Northeastern Atlantic and Mediterranean, with explanatory notes. Journal of Zoology, London 167: 15–29.

48. Carnevale G, Bannikov AF, Landini W, Sorbini C (2006) Volhynian (early Sarmatian sensu lato) fishes from Tsurevsky, North Caucasus, Russia. Journal of Paleontology 80: 684–699.

49. Oszczypko-Clowes M, ydek B (2012) Paleoecology of the Upper Eocene-Lower Oligocene Malcov Basin based on the calcareous nannofossils: a case study of the Luluchów section (Krynica zone, Magura Nappe, Polish outer Carpathians). Geologica Carpathica 63: 149–164.

50. Patterson C (1993) The Fossil Record 2, Chapman Hall, chapter Osteichthyes: Teleostei. pp. 621–656.

51. Nolf D (2003) Fish otoliths from the Santonian of the Pyrenean faunal province, and an overview of all otolith-documented North Atlantic Late Cretaceous teleosts. Bulletin de l’Institut Royal des Sciences Naturelles de Belgique, Sciences de la Terre 73: 155–173.

52. Caus E, Cornella A, Gallemi J, Gili E, Martinez R, et al. (1981) Field guide: Excursions to Conician-Maastrichtian of south central Pyrenees. Universidad Autonoma de Barcelona, Publicaciones de Geologia 13: 1–70.

53. Hottinger L, Drobne K, Caus E (1989) Late Cretaceous, larger, complex miliolids (Foraminifera) endemic in the Pyrenean faunal province. Facies 21: 99–134.

54. Reichenbacher B, Weidman M (1992) Fisch-otolithen aus der Oligo/Miozänen molasse der West-Schweiz und der Haute Savoie (Frankreich). Stuttgarter Beiträge zur Naturkunde, Serie B (Geologie und Paläontologie) 184: 1–83.

55. de Leeuw A, Mandic O, de Bruijn H, Marković Z, Reumer J, et al. (2011) Magnetostratigraphy and small mammals of the Late Oligocene Banovíci basin in NE Bosnnia and Herzegovina. Palaeogeography, Palaeoclimatology, Palaeoecology 340: 400–412.

56. Nolf D, Aguilera O (1998) Fish otoliths from the Cantaure formation (Early Miocene of Venezuela). Bulletin de l’Institut Royal des Sciences Naturelles de Belgique, Sciences de la Terre 68: 237–262.

57. Rey OT (1996) Estratigrafia de la península de Paraguaná, Venezuela. Revista de la Facultad de Ingegnería, Universidad Central de Venezuela 11: 35–45.

58. Grandstein FM, Ogg JG, Schmitz MD, Ogg GM (2012) The Geological Time Scale 2012. Amsterdam: Elsevier.

59. Suchard MA, Weiss RE, Sinsheimer JS (2001) Bayesian selection of continuous-time markov chain evolutionary models. Molecular Biology and Evolution 18: 1001–1013.

60. Xie W, Lewis PO, Fan Y, Kuo L, Chen MH (2011) Improving marginal likelihood estimation for Bayesian phylogenetic model selection. Systematic Biology 60: 150–160.

61. Fan Y, Wu R, Chen MH, Kuo L, Lewis PO (2011) Choosing among partition models in Bayesian phylogenetics. Molecular Biology and Evolution 28: 523–532.

62. Lartillot N, Philippe H (2006) Computing Bayes factors using thermodynamic integration. Systematic Biology 55: 195.

63. Baele G, Lemey P, Bedford T, Rambaut A, Suchard MA, et al. (2012) Improving the accuracy of demographic and molecular clock model comparison while accommodating phylogenetic uncertainty. Molecular Biology and Evolution 30: 2157–2167.

64. Baele G, Li WLS, Drummond AJ, Suchard MA, Lemey P (2013) Accurate model selection of relaxed molecular clocks in Bayesian phylogenetics. Molecular Biology and Evolution 30: 239–243.

65. Kass RE, Raftery AE (1995) Bayes factors. Journal of American Statistical Association 90: 773–795.

66. O’Brien JD, Minin VN, Suchard MA (2009) Learning to count: Robust estimates for labeled distances between molecular sequences. Molecular Biology and Evolution 26: 801–814.

67. Minin V, Suchard M (2008) Counting labeled transitions in continuous-time Markov models of evolution. Journal of Mathematical Biology 56: 391–412.

68. Raftery AE, Newton M, Satagopan J, Krivitsky P (2007) Estimating the integrated likelihood via posterior simulation using the harmonic mean identity. In: J M Bernardo MJB, Berger JO, editors, Bayesian Statistics. Oxford University Press, Oxford, pp. 1–45.

69. Cybis GB, Sinsheimer JS, Bedford T, Mather AE, Lemey P, et al. (2015) Assessing phenotypic correlation through the multivariate phylogenetic latent liability model. arXiv: 1406.3863v1.

70. Miller M, Pfeiffer W, Schwartz T. “Creating the CIPRES science gateway for inference of large phylogenetic trees”, in Proceedings of the Gateway Computing Environments Workshop (GCE), 14 Nov. 2010, New Orleans, LA, pp. 1–8.

71. Yang Z, Rannala B (2010) Bayesian species delimitation using multilocus sequence data. Proceedings of the National Academy of Sciences 107: 9264–9269.

72. Rannala B, Yang Z (2013) Improved reversible jump algorithms for Bayesian species delimitation. Genetics 195: 245–253.

73. Yang Z, Rannala B (2015) Unguided species delimitation using DNA sequence data from multiple loci. Molecular Biology and Evolution 32: DOI:10.1093/molbev/msu279.

74. Zhang C, Rannala B, Yang Z (2014) Bayesian species delimitation can be robust to guide-tree inference errors. Systematic Biology 63: 993–1004.

75. Venditti C, Meade A, Pagel M (2006) Detecting the node-density artifact in phylogeny reconstruction. Systematic Biology 55: 637–643.

76. Hugall AF, Lee MSY (2007) The likelihood node density e?ect and consequences for evolutionary studies of molecular rates. Evolution 61: 2293–2307.

77. Venditti C, Meade A, Pagel M (2008) Phylogenetic mixture models can reduce node-density artifacts. Systematic Biology 57: 286–293.

78. Ronquist F, Klopfstein S, Vilhelmsen L, Schulmeister S, Murray DL, et al. (2012) A total-evidence approach to dating with fossils, applied to the early radiation of the Hymenoptera. Systematic Biology 61: 973–999.

79. Pyron RA (2011) Divergence time estimation using fossils as terminal taxa and the origins of Lissamphibia. Systematic Biology 60: 466–481.

80. Bergmann C (1847) Ueber die Verhaltnisse der warmeokonomie der thiere zu ihrer grosse. Gottinger Studien 3: 595–708.

81. Vinarski M (2014) On the applicability of Bergmann’s rule to ectotherms: The state of the art. Biology Bulletin Reviews 4: 232–242.

82. Belk MC, Houston DD (2002) Bergmann’s rule in ectotherms: A test using freshwater fishes. American Naturalist 160: 803–808.

83. Wilson AB (2009) Fecundity selection predicts Bergmann’s rule in syngnathid fishes. Molecular Ecology 18: 1263–1272.

84. Blanchet S, Grenouillet G, Beauchard O, Tedesco PA, Leprieur F, et al. (2010) Non-native species disrupt the worldwide patterns of freshwater fish body size: implications for Bergmann’s rule. Ecology Letters 13: 421–431.

85. Fisher JAD, Frank KT, Leggett WC (2010) Global variation in marine fish body size and its role in biodiversity-ecosystem functioning. Marine ecology progress series 405: 1–13.

86. Rypel AL (2014) The cold-water connection: Bergmann’s rule in North American freshwater fishes. The American Naturalist 183: pp. 147–156.

87. Denys GP, Tedesco PA, Oberdorff T, Gaubert P (2015) Environmental correlates of body size distribution in Cyprinidae (Actinopterygians) depend on phylogenetic scale. Ecology of Freshwater Fish 24: DOI:10.1111/e?.12196.

